# Compression strategies for large-scale electrophysiology data

**DOI:** 10.1101/2023.05.22.541700

**Authors:** Alessio P. Buccino, Olivier Winter, David Bryant, David Feng, Karel Svoboda, Joshua H. Siegle

**Affiliations:** Allen Institute for Neural Dynamics, Seattle, WA; Champalimaud Foundation, Lisbon, Portugal; Independent Researcher, San Francisco, CA

**Keywords:** electrophysiology, data compression, spike sorting, high-density neural devices

## Abstract

With the rapid adoption of high-density electrode arrays for recording neural activity, electrophysiology data volumes within labs and across the field are growing at unprecedented rates. For example, a one-hour recording with a 384-channel Neuropixels probe generates over 80 GB of raw data. These large data volumes carry a high cost, especially if researchers plan to store and analyze their data in the cloud. Thus, there is a pressing need for strategies that can reduce the data footprint of each experiment. Here, we establish a set of benchmarks for comparing the performance of various compression algorithms on experimental and simulated recordings from Neuropixels 1.0 (NP1) and 2.0 (NP2) probes. For lossless compression, audio codecs (`FLAC` and `WavPack`) achieve compression ratios 6% higher for NP1 and 10% higher for NP2 than the best general-purpose codecs, at the expense of a slower decompression speed. For lossy compression, the `WavPack` algorithm in “hybrid mode” increases the compression ratio from 3.59 to 7.08 for NP1 and from 2.27 to 7.04 for NP2 (compressed file size of *∼*14% for both types of probes), without adverse effects on spike sorting accuracy or spike waveforms. Along with the tools we have developed to make compression easier to deploy, these results should encourage all electrophysiologists to apply compression as part of their standard analysis workflows.

## Introduction

In recent years, the field of extracellular electrophysiology has undergone a *big data* revolution. With the development, commercialization, and distribution of CMOS-based neural probes such as Neuropixels (Jun et al. 2017; Steinmetz et al. 2021),^1^ neuroscientists can now record brain function at unprecedented scales, accelerating progress towards a better understanding of the nervous system (Churchland et al. 2007).

Neuropixels 1.0 probes are capable of simultaneously sampling voltages from 384 electrodes at 30 kHz, increasing data throughput by an order of magnitude compared to previous methods. These probes have already been used to record spike trains from tens of thousands of neurons distributed across the entire brain (Steinmetz et al. 2019; Siegle et al. 2021; Chen et al. 2023). Recently developed SiNAPS probes (Angotzi et al. 2019; Boi et al. 2020) increase the number of simultaneously recorded electrodes per probe to 1024. And even higher electrode counts have been achieved in CMOS-based micro-electrode arrays (MEAs) for *in vitro* experiments, such as the 3-Brain HyperCAM Alpha (6144 channels)^2^ and the Maxwell Biosystems MaxTwo (26,400 channels).^3^ All these devices generate raw data at massive rates:

- Neuropixels 1.0: 30 kHz x 384 channels x 2 bytes/sample = *∼*80 GB/hour^4^
- SiNAPS 4-shank: 25 kHz x 1024 channels x 2 bytes/sample = *∼*185 GB/hour
- HyperCAM Alpha: 6 wells x 1024 channels x 20 kHz x 2 bytes/sample = *∼*885 GB/hour
- MaxTwo: 24 wells x 1024 channels x 20 kHz x 2 bytes/sample = *∼*3.5 TB/hour

As manufacturing shifts to even smaller CMOS processes, electrode counts will continue to grow (Buccino et al. 2022).

Sharing and processing data at this scale is nontrivial. The electrophysiology community has therefore begun migrating data and workloads to flexible, scalable pipelines that run in the public cloud (e.g., `DANDI` (Halchenko et al. 2022), `NeuroCAAS` (Abe et al. 2022), `DataJoint` (Yatsenko et al. 2015), and `Code Ocean` (Cheifet 2021)). Cloud platforms offer the ability to remotely launch standardized analysis environments that read from a single, immutable dataset, eliminating the need for individuals and labs to maintain local copies. However, cloud storage can be expensive. Consider a hypothetical example of a medium-sized experimental neuroscience laboratory using Neuropixels 1.0 probes. If each of seven lab members records an average of five one-hour sessions per month, the lab will produce *∼*33.5 TB of uncompressed data per year. Kept in the most performant tier of public cloud storage with an average cost of $0.023/GB per month,^5^ after five years data storage alone would incur an annual cost of *∼*$45,000. Effective compression may therefore be the deciding factor for whether a lab can afford scalable cloud technologies in their work.

Individual labs will not be the only ones facing mounting data storage costs. As of January 25^th^, 2023, all data associated with publications funded by the NIH must be made available through publicly-funded data repositories, many of which are backed by cloud technologies (National Institutes of Health 2021). Similar policies hold for work funded by the European Union (European Commission Directorate-General for Research and Innovation 2021). To avoid skyrocketing bills for archiving these datasets in perpetuity, funding agencies have a big incentive to determine the maximum amount of compression that can be reasonably applied to raw data.

Effective data compression is of paramount importance from both financial and environmental perspectives (because larger files require more energy to transfer and store). However, there has been little work investigating how compression algorithms perform on large-scale electrophysiology data. In this report, we describe a set of tools that make compression easier to deploy, and we use these tools to benchmark a wide array of compression strategies on both experimental and simulated electrophysiology data.

There are two general categories of compression, *lossless* and *lossy*, both of which are ubiquitous in everyday computing. Lossless compression reduces the size of a file without removing any information, meaning the original data will be perfectly intact following decompression. The final size of the compressed file depends on the randomness or redundancy of the data it contains (a file with more random/unpredictable values will be less compressible). Different lossless compressors employ different strategies for eliminating redundancy, but so far there has been no systematic comparison of how they interact with continuously sampled electrophysiology signals, which display high correlations across both space and time.

Lossy compression can further reduce file sizes by removing information that may be unnecessary for downstream workflows, but care must be taken to ensure that patterns of interest are preserved in the decompressed data. Lossy compression of image, video, and sound files is indispensable for the reliable functioning of the Internet, but it is rarely used in scientific applications. We believe that lossy compression of electrophysiology data will become increasingly critical as data volumes explode. But before this can be applied at scale, we must identify efficient compression algorithms that do not adversely affect the results of downstream analyses, especially spike sorting.

## Materials and Methods

### Compression framework

We chose `Zarr` as the software framework for running compression (Miles et al. 2020). The main reasons for this choice were:

- It is natively developed in `Python` and has a mature API.
- It stores data in a hierarchical directory structure that facilitates random access and streaming.
- It integrates a wide array of modern compression algorithms via the `numcodecs`^6^ package, which is developed and maintained by the same team. The `numcodecs` API makes it easy to implement and distribute additional codecs, which can be released as `Python` packages that are automatically registered in the `Zarr` framework.
- Neurodata Without Borders (Teeters et al. 2015; Rübel et al. 2022), a standard exchange format for neurophysiology data, can now read and write `.zarr` files (Tritt et al. 2019).^7^

We chose `Zarr` over `HDF5` (The HDF Group 2002), which is written in C++ and stores data in less cloud-friendly monolithic binary files. `Zarr` is also becoming the standard in bioimaging (see OME-NGFF – Moore et al. 2021).

To apply compression algorithms to large-scale electrophysiology data, we implemented a `Zarr` backend for `SpikeInterface` (Buccino et al. 2020), a Python package for building spike sorting pipelines (see Software extensions). `SpikeInterface` can read data from all widely used electrophysiology data formats (thanks to integration with the `neo` library) and, using the new backend, can output compressed `.zarr` files with a single line of code.

We also considered two electrophysiology-specific compression packages, but ultimately decided not to include them in our tests. The first, called `mtscomp`, is a Python library for lossless compression developed by the International Brain Laboratory. (International Brain Laboratory 2022)^8^. It implements delta filters and `zlib` compression, features that are also available via `Zarr`. We were thus able to test the same compression algorithm within `Zarr`. The second, called `MED` (Multiscale Electrophysiology Data) (Brinkmann et al. 2009; Stead & Halford 2016),^9^ specifies a custom data format that supports both lossless and lossy compression via predictive range-encoded differences, minimal bit encoding, variable frequency compression mode, and multiple derivative compression mode. While `MED` offers electrophysiologists a straightforward way to compress their data, the difficulty of integrating it with other open-source tools (such as `Zarr`) precluded us from testing it here.

### Benchmark datasets

We used several experimental datasets to test lossless compression algorithms, while both experimental and simulated datasets were used to assess the performance of lossy approaches. We focused on data from Neuropixels 1.0 (NP1) (Jun et al. 2017) and 2.0 (NP2) (Steinmetz et al. 2021) probes, given their broad adoption by the neuroscience community. However, we expect that our results and resources will be directly applicable to other types of recording devices.

### Experimental data

Half of the NP1 datasets are from the IBL Brain Wide Map (International Brain Laboratory 2022) (IBL in Table 1). We used the datasets in the `spikesorting/benchmark` folder on AWS, which includes four 20-minute (1200 s) recordings from various brain regions. Acquisition was performed with `SpikeGLX` software (Karsh, n.d.). The other half of the NP1 datasets and all NP2 datasets were pilot sessions acquired by the Allen Institute of Neural Dynamics using `Open Ephys` software (Siegle et al. 2017) and targeting diverse brain regions (AIND in Table 1). Recording durations ranged from 732 s to 2430 s.

**Table 1:** Overview of the experimental and simulated datasets

| Session | Source | Type | Probe | Duration (s) | Acquisition software | Band | Gain | LSB |
| --- | --- | --- | --- | --- | --- | --- | --- | --- |
| CSHZAD026_2020-09-04_probe00 | IBL | exp | NP1 | 1200 | SpikeGLX | spike | 2.34 | 1 |
| CSHZAD029_2020-09-09_probe00 | IBL | exp | NP1 | 1200 | SpikeGLX | spike | 2.34 | 1 |
| SWC054_2020-10-05_probe00 | IBL | exp | NP1 | 1200 | SpikeGLX | spike | 2.34 | 1 |
| SWC054_2020-10-05_probe01 | IBL | exp | NP1 | 1200 | SpikeGLX | spike | 2.34 | 1 |
| 625749_2022-08-03_15-15-06_ProbeA | AIND | exp | NP1 | 1817 | Open Ephys | spike | 0.195 | 12 |
| 634568_2022-08-05_15-59-46_ProbeA | AIND | exp | NP1 | 1766 | Open Ephys | spike | 0.195 | 12 |
| 634569_2022-08-09_16-14-38_ProbeA | AIND | exp | NP1 | 1741 | Open Ephys | spike | 0.195 | 12 |
| 634571_2022-08-04_14-27-05_ProbeA | AIND | exp | NP1 | 1815 | Open Ephys | spike | 0.195 | 12 |
| 595262_2022-02-21_15-18-07_ProbeA | AIND | exp | NP2 | 732 | Open Ephys | wide | 0.195 | 3 |
| 602454_2022-03-22_16-30-03_ProbeB | AIND | exp | NP2 | 901 | Open Ephys | wide | 0.195 | 3 |
| 612962_2022-04-13_19-18-04_ProbeB | AIND | exp | NP2 | 1030 | Open Ephys | wide | 0.195 | 3 |
| 612962_2022-04-14_17-17-10_ProbeC | AIND | exp | NP2 | 1055 | Open Ephys | wide | 0.195 | 3 |
| 618197_2022-06-21_14-08-06_ProbeC | AIND | exp | NP2 | 745 | Open Ephys | wide | 0.195 | 3 |
| 618318_2022-04-13_14-59-07_ProbeB | AIND | exp | NP2 | 949 | Open Ephys | wide | 0.195 | 3 |
| 618384_2022-04-14_15-11-00_ProbeB | AIND | exp | NP2 | 1273 | Open Ephys | wide | 0.195 | 3 |
| 621362_2022-07-14_11-19-36_ProbeA | AIND | exp | NP2 | 2430 | Open Ephys | wide | 0.195 | 3 |
| MEAREC_NP1 | MEAREC | sim | NP1 | 600 | MEAREC | spike | 0.195 | 12 |
| MEAREC_NP2 | MEAREC | sim | NP2 | 600 | MEAREC | spike | 0.195 | 3 |

There are several key differences between NP1 and NP2 datasets that are expected to impact compression performance. NP1 devices include an analog 300-Hz high-pass filter associated with the AP (*action potential*) stream, which is sampled at 30 kHz. The same set of channels is also low-pass filtered at 1 kHz and sampled at 2.5 kHz to produce the *LF* (*local field*) stream (Dutta et al. 2019). We only tested compression performance on the *AP* stream. NP2 devices do not include comparable hardware filters and instead produce a single broadband stream. In addition, although both NP1 and NP2 data are saved to disk as 16-bit integers, the NP1 device includes analog-to-digital converters (ADCs) with 10-bit resolution, while NP2 includes ADCs with 14-bit resolution.^10^ The higher-bit-depth ADCs of NP2 are used both to record at 4*×* higher resolution (0.585 µV/sample for NP2 vs. 2.34 µV/sample for NP1) and to accommodate the dynamic range of the wide-band signals.

We also considered the effects of the acquisition software, which can apply different scaling factors to the data prior to saving. `SpikeGLX` outputs the raw values of the Neuropixels ADCs, which must be scaled to account for the hardware amplifier gain prior to analysis. In this case, the data has a least significant bit (LSB) of 1, regardless of which type of device was used. `Open Ephys` scales the data to have a final gain of 0.195 µV/sample, regardless of the hardware gain settings. This decision was made to minimize the effects of rounding errors if online processing steps, such as band-pass filtering, are applied before saving the data. However, if the raw Neuropixels data was written, as it was in all the example datasets we analyzed, this rescaling makes the LSB equal to 12 for NP1 and 3 for NP2. Prior to compression, we rescaled the `Open Ephys` data to have an LSB of 1 by first removing each channel’s median (since scaling could introduce rounding errors) and dividing by either 12 or 3.

### Simulated data

We used the `MEArec` software (Buccino & Einevoll 2021) to simulate voltage traces that resemble the Neuropixels 1.0 and 2.0 data as recorded by the `Open Ephys Neuropix-PXI` plugin (same acquisition device as the experimental data from the AIND sources, see Table 1). We refer to these datasets as ground truth (GT), since the underlying spiking activity is known. We simulated one dataset per probe type with the following characteristics:

- data type: `int16`
- gain: 0.195 µV/sample
- number of ground-truth cells: 100
- noise level: 10 µV additive spatially correlated noise

The LSB was set to 12 for NP1 and 3 for NP2 to match the experimental settings.

Ground-truth recordings allow us to test whether lossy strategies significantly affect waveform features (in this case we can use the ground-truth spiking activity to compute waveforms from the lossy recordings) and also to quantitatively benchmark spike sorting outcomes. For the latter case, our goal is not to quantify the overall accuracy of a particular spike sorter but rather to measure how sorting results change with increasing levels of lossiness. For this reason, the simulated datasets are relatively easy to sort, with only 100 ground truth units, no additional drift, and no burst-related waveform attenuation.

### Lossless compression

#### Codecs

We benchmarked most of the codecs in the `Zarr`-`numcodecs` library. The `BLOSC` filter (The Blosc Development Team, n.d.) is a high-performance meta-compressor optimized for binary data. It includes four codecs—`LZ4`, `LZ4HC` (Collet, n.d.), `ZLIB` (Gailly & Adler, n.d.), `ZSTD` (Facebook, n.d.)—and two shuffling options (bit, byte).^11^ These codecs are referred to as *blosc* compressor type. In addition to the `BLOSC` interface, we tested other codecs that are directly wrapped in `numcodecs` (including some also available through the `BLOSC` filter): `GZIP` (Gailly & Adler, n.d.), `LZMA` (7zip) (Pavlov, n.d.), `LZ4`, `ZLIB`, and `ZSTD`. These *numcodecs* compressor types were also tested with the byte shuffling option, available with the `numcodecs.Shuffle` filter.

Both *blosc* and *numcodecs* compressor types are general-purpose compressors, i.e., they make no assumption about the structure of the data. Electrophysiology data have many features in common with audio signals: they are both time series that are sampled at a few tens of kHz (usually 30 kHz for electrophysiology and 44.1 kHz for audio), they have similar frequency content, and they have channels that may be strongly correlated (e.g., left/right stereo signals or nearby electrodes in neural probes). Given these similarities, we also tested the performance of lossless compressors designed for audio signals, including `FLAC` (xiph.org Foundation et al., n.d.) and `WavPack` (Bryant, n.d.). Lossless audio codecs like `FLAC` and `WavPack` operate with two basic steps. The first step attempts to model the characteristics of the data to produce a prediction of the sampled values. Then, for each sample, this prediction is subtracted from the actual sample to generate an error value, sometimes referred to as the “residual.” This step is called *decorrelation* because its purpose is to remove the correlation (or redundancy) that normally exists in sampled signals. The correlation is generally highest in consecutive samples in the same channel; however, there is often correlation *between* channels that can also be utilized. Once the samples are represented as residuals, the next step is entropy encoding, which refers to storing the samples with the minimum number of bits. `FLAC` uses the simplest of these entropy encoding methods (Golomb-Rice codes). `WavPack` uses a more complex scheme that provides better compression but is more computationally costly. Table 2 lists all the codecs that have been tested in this study.

**Table 2:** Overview of the lossless compression codecs, shuffling options, channel block sizes, and compression levels mapped to codec-specific parameters. For channel block size, a value of -1 indicates the array was flattened prior to compression.

| compressor | compressor type | shuffling options | channel block size | (low, medium, high) levels |
| --- | --- | --- | --- | --- |
| blosc-lz4 | blosc | (no, byte, bit) | -1 | (1, 5, 9) |
| blosc-lz4hc | blosc | (no, byte, bit) | -1 | (1, 5, 9) |
| blosc-zlib | blosc | (no, byte, bit) | -1 | (1, 5, 9) |
| blosc-zstd | blosc | (no, byte, bit) | -1 | (1, 5, 9) |
| gzip | numcodecs | (no, byte) | -1 | (1, 5, 9) |
| lzma | numcodecs | (no, byte) | -1 | (1, 5, 9) |
| lz4 | numcodecs | (no, byte) | -1 | (9, 5, 1) |
| zlib | numcodecs | (no, byte) | -1 | (1, 5, 9) |
| zstd | numcodecs | (no, byte) | -1 | (1, 11, 22) |
| flac | audio | (no) | (-1, 2) | (1, 5, 8) |
| wavpack | audio | (no) | (384) | (1, 2, 3) |

#### Chunk size

The `Zarr` framework supports the storage (and compression) of N-dimensional arrays in chunks. Chunks are stored in separate files that are compressed independently, which is critical for performant, parallel encoding and decoding. In the case of extracellular electrophysiology, the input data is two-dimensional, with a shape of (*n*_*samples*, *n*_*channels*). For Neuropixels data, *n*_*channels* is 384, since this is the number of channels that can be recorded simultaneously. For all codecs, we explored three different chunk sizes in time: 0.1 s, 1 s, and 10 s (which correspond to 3,000, 30,000, and 300,000 samples per chunk). For general-purpose compressors (compressor types *blosc* and *numcodecs*), which flatten an N-dimensional array before compression, we only chunked in time, so that the shape of each chunk is (*n*_*samples*_*chunk*, *n*_*channels*) (denoted as -1 in the *channel block size* column of Table 2). For the `FLAC` compressor, whose `Python` implementation^12^ supports both mono and stereo (2-channel) signals, we utilized either flattening before compression (*channel block size* = -1) or blocks with 2 channels (to mimic stereo signals). The `WavPack` codec supports any number of channels, and it internally constructs streams of stereo pairs (with a final mono stream to handle odd channel counts) prior to compression. Therefore, for `WavPack` we used a chunk size of (*n*_*samples*_*chunk*, *n*_*channels*).

#### Compression level

All compressors listed in Table 2 allow different levels of compression. Higher levels achieve better compression but are usually slower in compression and decompression speeds. Because each codec may use different numeric values to specify compression level, we defined three compression levels for each codec evenly distributed over the available range. The codec-specific compression levels corresponding to *low*, *medium*, and *high* are listed in the right-most column of Table 2.

#### Delta filters and pre-processing

A common pre-compression operation is to apply a *delta* filter, which encodes data as the difference between adjacent values to reduce the dynamic range of the signals. While the `numcodecs` library provides an implementation of such a filter (`numcodecs.Delta`), this implementation is targeted to one-dimensional input data, and it flattens any multi-dimensional input arrays. In the case of electrophysiology data, the input is two-dimensional (*n*_*samples*, *n*_*channels*), and it is therefore convenient to apply a delta filter across specific axes. To do so, we implemented a `Delta2D` codec (see Software extensions) which allowed us to apply a delta filter across time, across channels (or *space*), or across both directions sequentially.

Electrophysiology data contain both local field potentials (<300 Hz) and spiking activity (>300 Hz). As mentioned in the Experimental data section, NP1 probes output two separate streams for these two signals (*LF* and *AP*), while NP2 probes do not apply hardware filters and output a single wide-band stream. We therefore investigated whether pre-processing the input data affects compression performance by applying either a high-pass filter (cutoff frequency at 300 Hz) or band-pass filters (with cutoffs at 300-6000 Hz and 300-15000 Hz), which is a common alternative for pre-processing electrophysiology data. We also tested the application of an anti-aliasing band-pass filter, with cutoffs at 0.5-15000 Hz. In all cases, we used 5*^th^*-order Butterworth filters implemented in the `SpikeInterface.preprocessing` module (highpass^_^filter() and bandpass^_^filter() functions) prior to compression.

### Lossy compression

We investigated two lossy compression strategies: bit truncation and the hybrid mode of the WavPack compressor.

#### Bit truncation

As reported in previous sections, NP1 and NP2 data have a resolution of 2.34 µV/sample and 0.585 µV/sample, respectively. Noise levels for Neuropixels probes are reported to be around 5.9 µV*_rms_* for NP1 (Jun et al. 2017) and slightly higher (7.4 µV*_rms_*) for NP2 (Steinmetz et al. 2021).^13^ In contrast, spike amplitudes of neurons that are usually considered well-isolated have amplitudes greater than 30 µV, implying that the voltage signal is over-sampled.

The bit truncation strategy reduces the resolution of the original data by “right-shifting” each sample. A right shift (i.e., bit truncation) of *n* bits is equivalent to dividing the sampled values, and consequently decreasing the resolution, by 2*^n^*. As an example, a bit truncation of NP1 data by 2 bits will result in a signal resolution of: 2.34 µV/sample *·*2^2°^= 9.36 µV/sample. Note that the NP2 probe has a resolution four times larger than NP1, hence a 2-bit truncation of NP2 data will match the 2.34 µV/sample resolution of NP1 probes. The bit truncation operation should increase the compression performance because it effectively adds *n* 0s to replace the truncated bits, reducing the amount of information that each sample carries.

Bit truncation is implemented with the `numcodecs.FixedScaleOffset` as a pre-compression filter, using a scale of 1/2*^n^*. This strategy was combined with the *blosc-zstd* compressor, which performed best among the general-compressor codecs. We tested truncations of up to 7 bits for both probe types.

#### WavPack Hybrid mode

The `WavPack` codec includes a hybrid (lossy) mode in addition to the lossless mode.^14^ To implement the hybrid compression mode, `WavPack`’s entropy encoder provides an optional “error limit” parameter which can be finely and continuously controlled by the characteristics of the input samples to target a given bit rate in units of bits per sample (*bps*). Because this error limit changes slowly, transients are more accurately encoded than with schemes that are restricted to a fixed number of bits per sample (like Adaptive Differential Pulse Code Modulation, or ADPCM). This is not only useful for the audio compression applications for which it was optimized, but also for the current application which relies on accurate reproduction of fast transients (i.e., spikes). We tested *bps* values of 6, 5, 4, 3.5, 3, 2.5, 2.25 (from least to most aggressive lossy compression). Note that 2.25 is the minimum value accepted by the `WavPack` hybrid mode.

### Evaluation and benchmarks

We used three metrics to quantify compression performance:

- **Compression Ratio (CR)**, computed by dividing the number of bytes in the original data file by the number of bytes written to disk after compression. The higher the compression ratio, the lower disk space the compressed data will occupy. For example, a CR of 2 implies that the compressed data occupies half of the disk space (50%) compared to uncompressed data, a CR of 4 occupies 25% of the disk space, and so on.
- **Compression speed** (in units of *x*RT, or “times real-time”), computed by dividing the time needed to compress an entire recording divided by the original recording duration (see *duration* in Table 1).
- **Decompression speed** (in *x*RT), computed by dividing the time needed to decompress all channels (384) in a 10 s time range by 10.

The three metrics should not be weighted equally when evaluating overall performance. Compression ratio directly relates to storage size and hence storage costs. Compression speed is potentially less important since, ideally, a given dataset will only need to be compressed once. Decompression speed is more important because decompression will be performed whenever the raw data must be accessed for analysis or visualization.

For lossy approaches, some information is irreversibly lost during compression. To quantify this loss of information, we additionally compute the root-mean-square error (RMSE) between the original and the decompressed lossy traces over 10 s (after applying a 300-6000 Hz band-pass filter). Since even small RMSE could affect downstream analysis, we also evaluate whether the compression (1) affects spike sorting results or (2) significantly changes spike waveforms. The latter requirement is important because features derived from the average extracellular waveform, such as peak-to-valley duration, full-width half maximum duration, and peak-to-trough ratio, can provide information about the cell type identity of recorded neurons (Buccino et al. 2018; Jia et al. 2019).

We used both simulated and experimental data to examine the effects of lossy compression on spike sorting. In all cases, we used `Kilosort2.5` (Pachitariu et al. 2016; Pachitariu et al. 2023) with default parameters after preprocessing with a band-pass filter and common median reference (the latter was only applied to experimental data, since simulations do not have common noise sources by construction). After spike sorting, we applied a minimal post-processing step to remove duplicated units sharing more than 95% of spikes with other units, keeping the units with more spikes. For simulated recordings, we compute the accuracy, precision, and recall (see Buccino et al. 2020 for details) of the spike sorting outputs and compared them to the values obtained on the original data. For experimental data, due to the lack of ground truth spike times, we counted the number of passing and failing units following automatic curation based on three spike sorting quality metrics (Siegle et al. 2021; inter-spike-interval violations ratio < 0.1, presence ratio > 0.9, and amplitude cutoff < 0.1). We report the fraction of passing and failing units with respect to the counts obtained by spike sorting the original uncompressed data. In addition, we also compare the full spike trains of all units detected after lossy compression to matched units in the lossless dataset.

We used simulated datasets to evaluate the impact of lossy compression on spike shape, since they allowed us to compare waveform features to the ground-truth average waveforms. We computed the three aforementioned metrics (peak-to-valley duration, full-width half maximum duration, and peak-to-trough ratio) both on the *main* channel with the largest amplitude for each unit (i.e., the unit main channel) and on a *peripheral* channel 60 µm away. This was done to ensure that not only are peak waveforms preserved, but features on peripheral channels are also well estimated. Note that waveform features computed using multiple channels, or 2D features, have been shown to improve cell type classification (Jia et al. 2019).

### Statistical analysis

For comparisons between two samples, we used the non-parametric Mann-Whitney U test; for population tests (more than two samples), we used the non-parametric Kruskal-Wallis test. Post-hoc tests were run with Conover’s test, with Holm’s correction for p-values. Tests were performed using `scipy` (Virtanen et al. 2020) and `scikit-posthocs` (Terpilowski 2019). In addition to reporting p-values for two-sample tests, we also report the Cohen’s d coefficient as a proxy for the *effect size*. In figures, we denote p-values using the following symbols:

- **\***: 0.001 *≤ p* < 0.01
- **\*\***: 10^−4^ *≤ p* < 0.001
- **\*\*\***: *p* < 10^−4^.

The values of compression metrics are always reported as the median *±* standard deviation of the associated distribution. We used the median due to its lower sensitivity to outliers. The shaded diamonds in all figures represent outliers as defined in (Hofmann et al. 2017).

## Results

### Lossless compression

We ran a total of 6300 independent compression jobs using a 16-CPU AWS EC2 instance with 128 GB of RAM, coordinated via the Code Ocean compute platform. In all cases, compression ran with 16 parallel threads, while decompression was not explicitly parallelized.^15^

#### LSB correction improves compression performance

As mentioned in the Experimental data section, the Neuropixels data recorded by `Open Ephys` (in our case from the `AIND` data source) do not have a unitary least significant bit (LSB = 12 for NP1, LSB = 3 for NP2). In principle, dividing the samples by the LSB value would reduce the data range (and possibly improve compression) without affecting data integrity. We denote this process as *LSB correction*. Compression ratio is significantly higher after LSB correction, with a large effect size (Figure 1a; *p* < 10^−10^, effect size = 0.99). LSB correction decreases compression speed, likely due to the additional computational load of the median subtraction and LSB division (Figure 1b; *p* < 10^−10^, effect size = 0.42), while slightly increasing decompression speed (Figure 1c; *p* < 0.01, effect size = 0.08). Note that in this case, the distributions include observations for both probe types (1800 NP1 and 3600 NP2). Due to the large improvement in compression ratio, we continued the analysis with LSB correction. We advise users to do so as well when using `Open Ephys` or other acquisition software that saves data with an LSB > 1, assuming that no other processing steps (such as a band-pass filter) have been applied (see the Software extensions section for more details).

**Figure 1:**
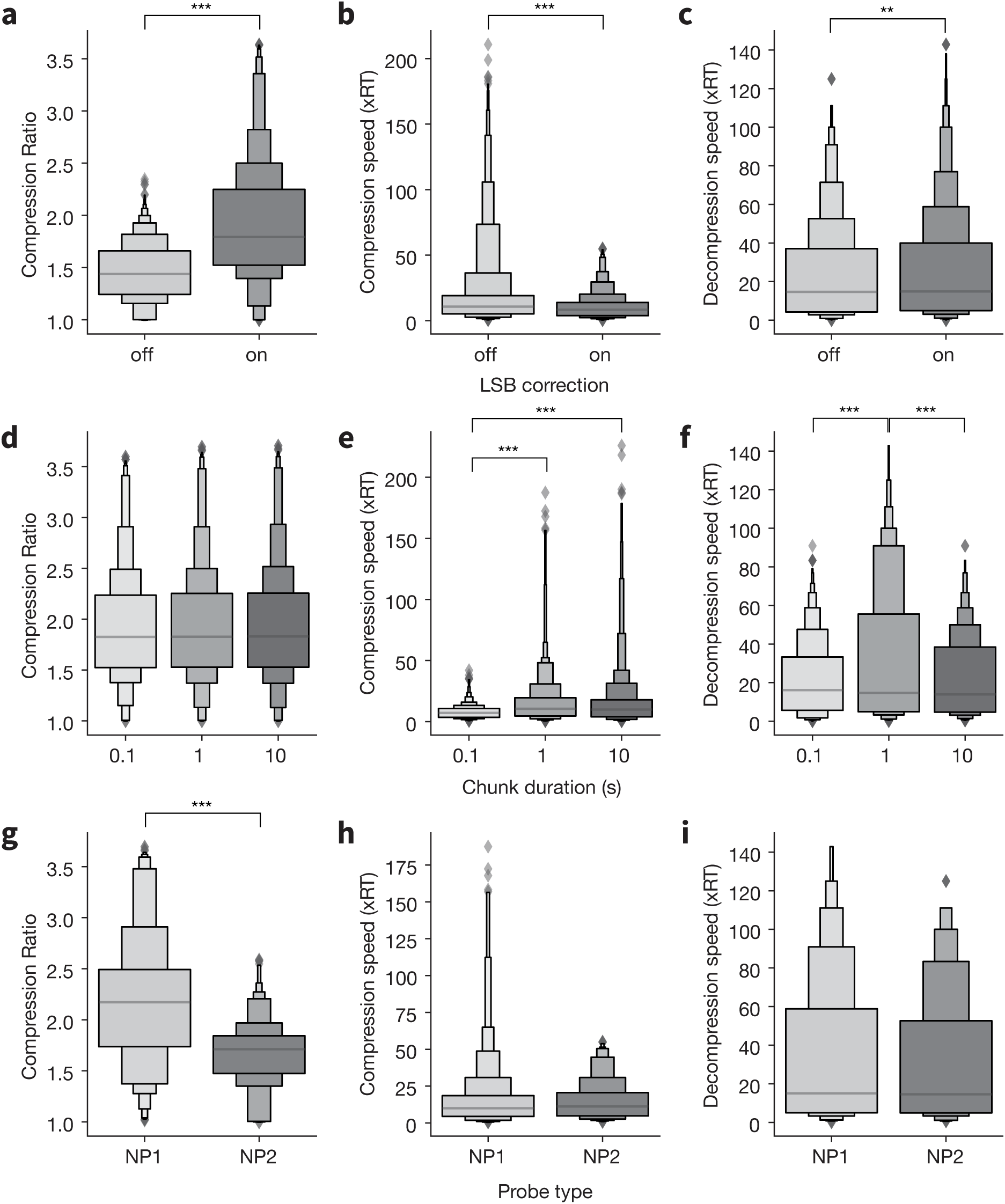
Effect of least significant bit (LSB) correction, chunk duration, and probe type on compression metrics. **a-c**, Effect of LSB correction on compression metrics (`Open Ephys` datasets only, *N* = 2700 compression jobs per condition, including 900 with NP1 data and 1800 with NP2 data). **d-f**, Effect of chunk duration on compression metrics. Data points without LSB correction are excluded (*N* = 1200 compression jobs per chunk duration). **g-i**, Effect of probe type on compression metrics (*N* = 600 compression jobs per probe type). Shaded diamonds represent outliers as defined in (Hofmann et al. 2017). **\*\***: 10^−4^ *≤ p* < 0.001, **\*\*\***: *p* < 10^−4^.

#### Chunk size affects compression speed, but not compression ratio

We analyzed compression metrics as a function of chunk duration, i.e., the size of each block fed into the compression algorithm (0.1 s, 1 s, and 10 s). We did not detect differences in compression ratio (Figure 1d), while compression speed was significantly slower for chunks of 0.1 s (Figure 1e; 0.1 *vs* 1: *p* < 10^−10^, effect size = 0.57; 1 *vs* 10: *p* < 10^−10^, effect size = 0.49). Chunks of 1 s also resulted in higher decompression speeds with respect to the other chunk sizes (Figure 1f; 0.1 *vs* 1: *p* < 10^−5^, effect size = 0.42; 1 *vs* 10: *p* < 10^−4^, effect size = 0.37). Given this significant increase in decompression speeds, and provided that 1 s chunk sizes are commonly used for parallel processing (e.g., they are the default in `SpikeInterface`), we restricted further analysis to chunk durations of 1 s (1200 total compression options).

#### Neuropixels 1.0 data is more compressible than 2.0 data

As described in the section, NP1 and NP2 devices have different properties that could impact the compressibility of the data they generate: NP1 probes split each channel into separate *AP* and *LF* bands, both of which are digitized with 10-bit ADCs. NP2 probes acquire broadband signals with 14-bit ADCs. We found that compression ratio was significantly higher for NP1 than NP2 (Figure 1g; *p* < 10^−10^, effect size = 1), while compression and decompression speeds were similar for both probe types (Figure 1h-i). This confirms the higher compressibility of NP1 data, with its more limited frequency content and lower ADC bit depth (see Table 1). Therefore, we analyze the results from the two probes separately in the following sections.

#### Comparison of nine general-purpose codecs

Figure 2 shows compression metrics for all the general-purpose codecs, applied separately to NP1 and NP2 data. The same trends can be observed for both probe types: `LZMA` produces the highest average compression ratio, immediately followed by `zstd`, whose *blosc* implementation (`blosc-zstd`) appears to outperform the *numcodecs* version (`zstd`). Although `LZMA` achieved a higher compression ratio with respect to `blosc-zstd` (NP1 - `LZMA`: 2.52*±*0.31, `blosc-zstd`: 2.38*±*0.36; NP2 - `LZMA`: 1.85*±*0.07, `blosc-zstd`: 1.76*±*0.15), its decompression speed is very slow (NP1 - `LZMA`: 0.89*±*0.41, `blosc-zstd`: 71.42*±*32.89; NP2 - `LZMA`: 0.81*±*0.94, `blosc-zstd`: 51.32*±*30.99). Note that, in general, *blosc* codecs achieve a much higher decompression speed than codecs directly implemented in *numcodecs*. As mentioned in the Evaluation and benchmarks section, this is probably due to the multi-threaded implementation of the `BLOSC` meta-compressor.

**Figure 2:**
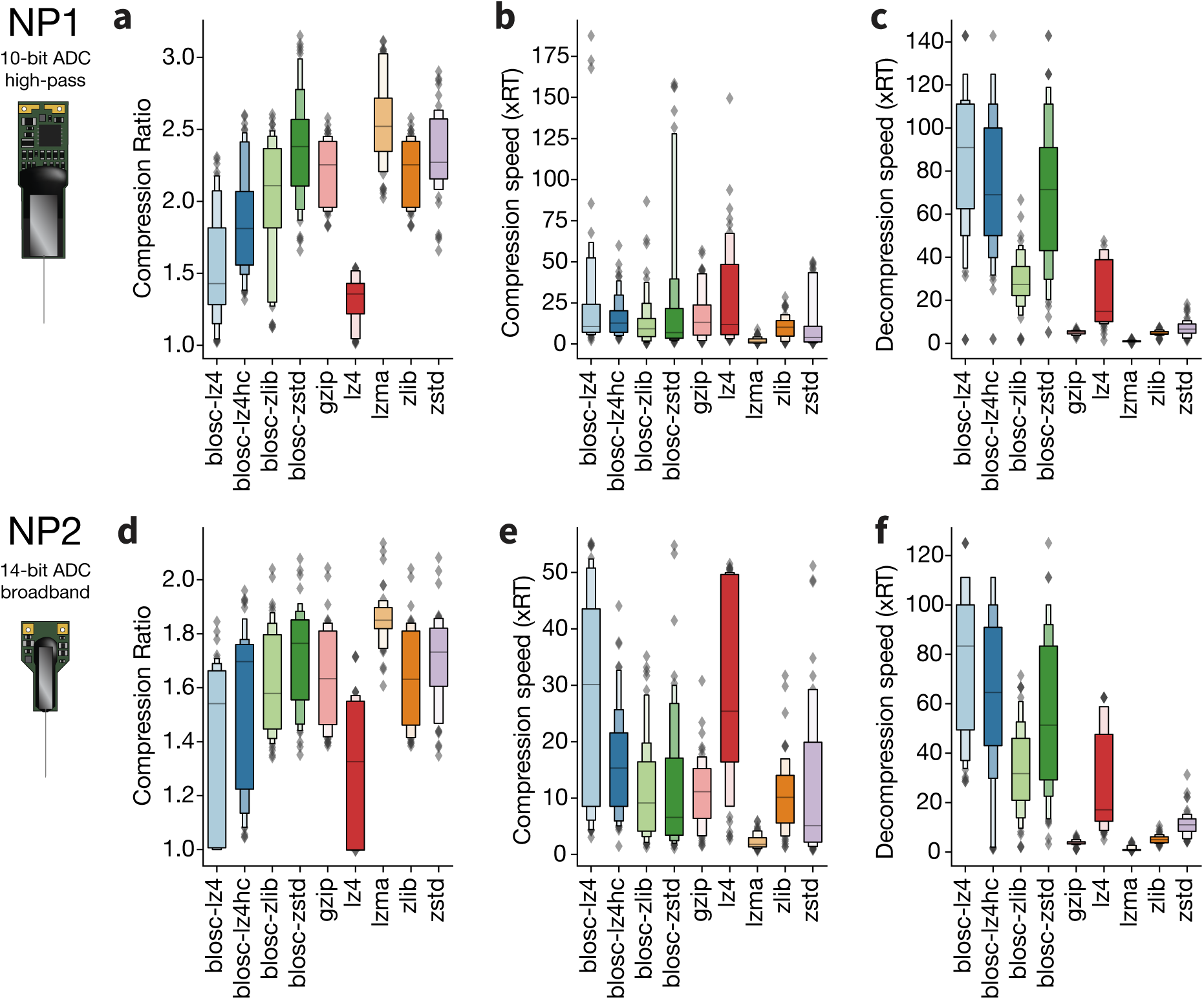
Comparing performance of general-purpose lossless codecs. Distributions of compression ratios (**a** and **d**), compression speeds (**b** and **e**), and decompression speeds (**c** and **f**) for nine different lossless codecs (*N* = 72 jobs for *blosc* compressors, *N* = 48 jobs for *numcodecs* compressors). Top row is for NP1, bottom row is for NP2.

The distributions shown in Figure 2 aggregate data points from all shuffling options and compression levels. Figure 3 breaks down the results based on different shuffling options. Shuffling improves compression metrics, especially compression ratio, for most compressors and both probe types (with the exception of `LZMA` for NP1, for which the highest compression ratio is achieved when no shuffling is applied). For *blosc* codecs, bit shuffling is also available and yields better compression metrics (except for the `blosc-zlib` codec, which does not behave well with bit shuffling for *medium* and *high* compression levels, see Figure S1).

**Figure 3:**
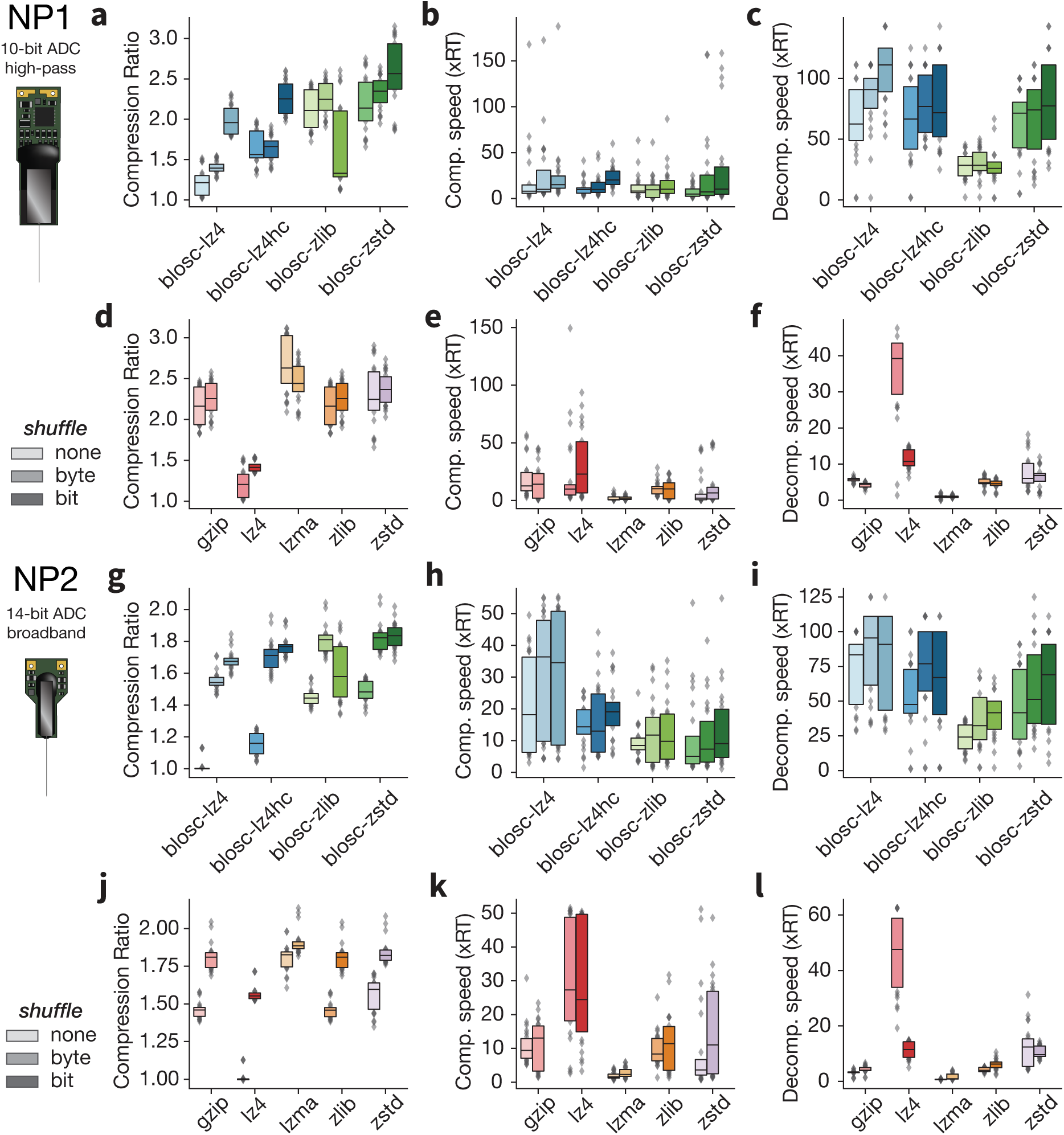
Effect of shuffling on compression performance. Distributions of compression ratios, compression speeds, and decompression speeds for different pre-shuffling options. *N* = 24 compression jobs per condition. The upper half (**a-f**) is for NP1, the lower half is for NP2 (**g-l**). The *blosc* codecs (**a-c** and **g-i**) support bit and byte shuffling, while `Zarr`’s built-in *numcodecs* (**d-f** and **j-l**) only support byte shuffling.

Figure 4 shows the effect of varying compression level. As expected, as compression level increases from *low* to *high*, one observes a higher compression ratio and lower compression/decompression speed (with the exception of `blosc-zlib`, whose average compression ratio decreases between the *low* and *medium* settings). Figure S2, which displays the compression ratio and decompression speed for each compressor depending on both the compression level for the shuffling option, confirms the same trends and conclusions that we observed when looking at the shuffling and compression level parameters separately.

**Figure 4:**
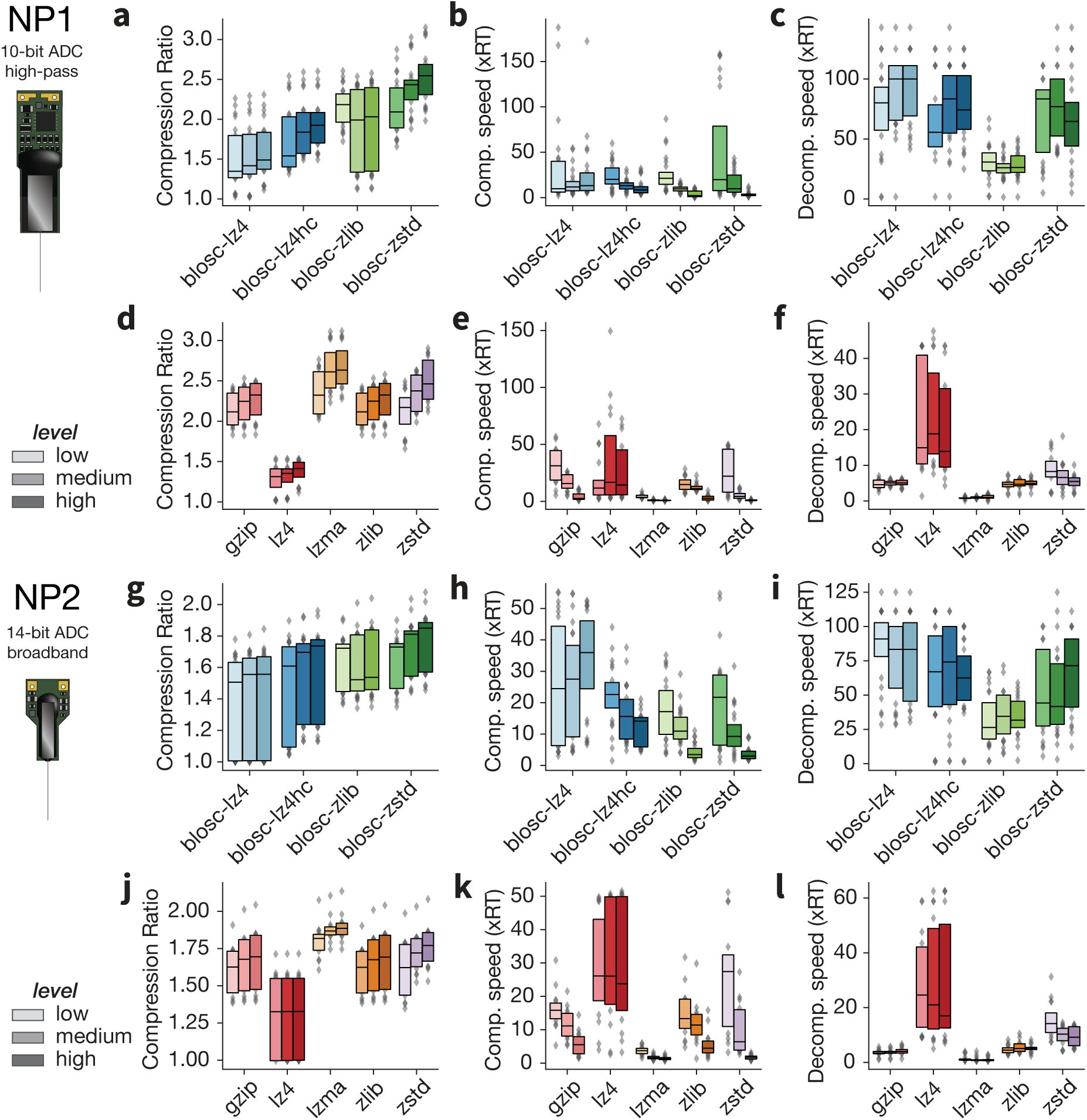
Effect of compression level on performance. Distributions of compression ratios, compression speeds, and decompression speeds for different compression levels. *N* = 24 jobs for each *blosc* codec (**a-c** and **g-i**), *N* = 18 jobs for *numcodecs* (**d-f** and **j-l**). The upper half (**a-f**) is for NP1, the lower half is for NP2 (**g-l**)

#### Audio codecs achieve a higher compression ratio than general-purpose codecs

Next, we investigated compression using *audio* codecs, namely `FLAC` and `WavPack`. Before directly comparing the two, we observed how channel chunking options affected `FLAC` performance. By default, the `FLAC` codec flattens the 384 Neuropixels channels into a single channel before compression (*channel block size* = -1). Enabling the option to group channels as stereo pairs (*channel block size* = 2) largely improves compression ratio (Figure S3a; *p* < 10^−3^, effect size: 0.81) at the expense of compression speed (Figure S3b; *p* < 10^−10^, effect size: 1.75) and decompression speed (Figure S3c; *p* < 10^−10^, effect size: 7.7). Since compression ratio is our main metric of interest, we selected *channel block size* = 2 for further comparisons (see the Chunk size for more details).

The `FLAC` and `WavPack` codecs yield similar compression ratios, with slightly higher values for `WavPack` for NP1 (Figure 5a), but `FLAC` for NP2 (Figure 5d). While both codecs see an improvement in compression ratio when transitioning from *low* to *medium* compression levels, the increase between *medium* and *high* is almost null (< 0.01). However, there is a sharp decrease in compression speed at *high* compression (Figure 5b,e), and decompression speed is also reduced for `WavPack` (Figure 5c,f). Given these results, we used the *medium* compression level for both audio codecs for all additional comparisons.

**Figure 5:**
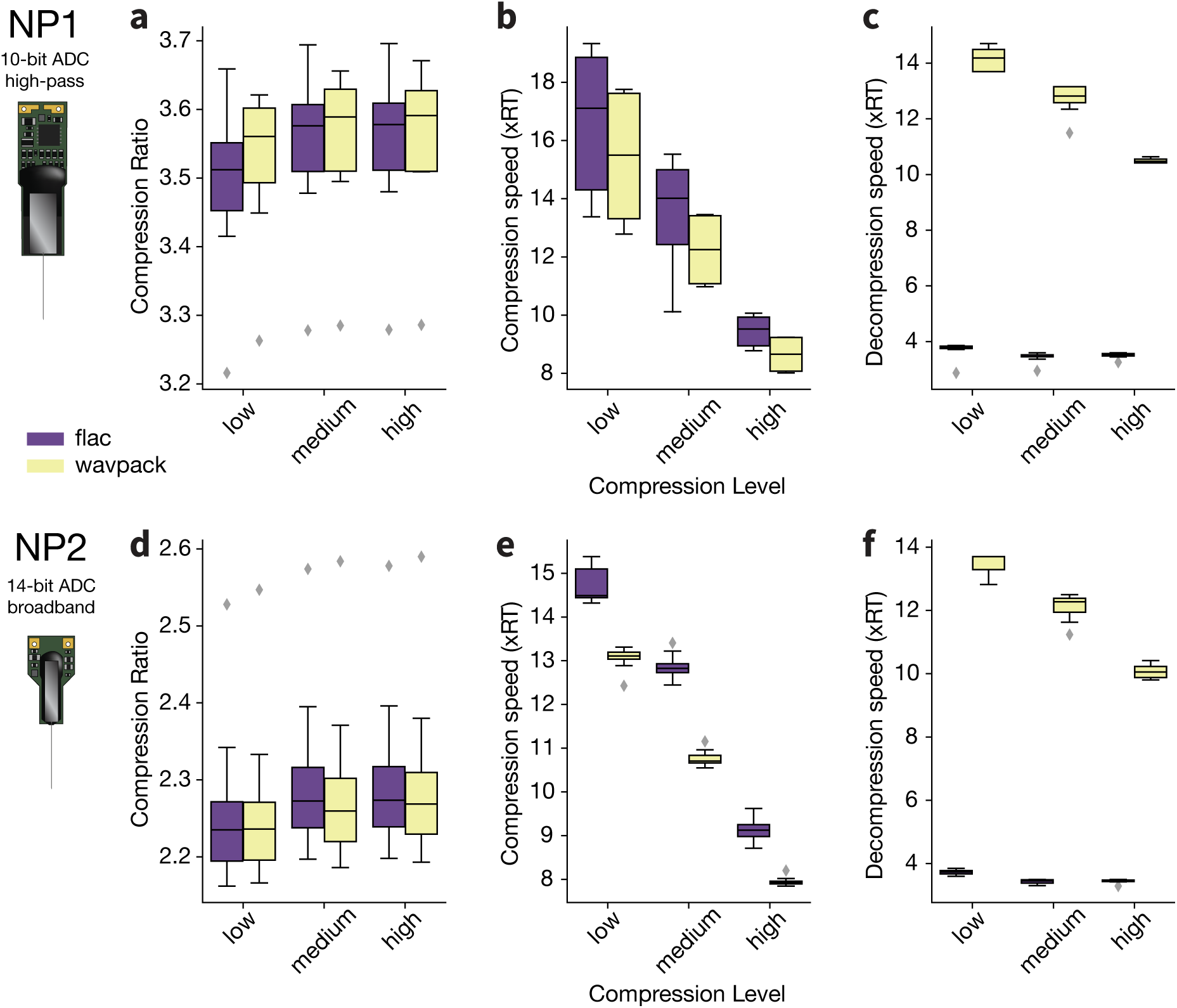
Performance of audio codecs. Compression ratios, compression speeds, and decompression speeds for the `FLAC` (purple) and `WavPack` (yellow) codecs, at three different compression levels (low, medium, high). The top row (**a-c**) is for NP1, the bottom row (**d-f**) is for NP2. *N* = 8 compression jobs for all distributions.

In Figure 6, we show compression metrics for the two `Zarr` codecs that achieved the best performance, `blosc-zstd` and `LZMA`, alongside metrics for the two audio codecs, `FLAC` and `WavPack`. For `LZMA`, we selected no shuffling for NP1 and byte shuffling for NP2, since these options achieved the highest compression ratio for the respective probe types. We selected the *high* compression level for `Zarr` compressors and *medium* for *audio* compressors, resulting in eight data points for each probe. Audio codecs clearly outperform general-purpose codecs in terms of compression ratio, with `WavPack` achieving a CR of 3.59*±*0.12 for NP1 (file size *∼*27.9%) and 2.26*±*0.13 for NP2 (file size *∼*44.2%). The compression ratio of `FLAC` is almost identical to `WavPack` (NP1: 3.58*±*0.13; NP2: 2.27*±*0.12), but `WavPack` is faster at decompression (`FLAC`: 3.51*±*0.21 *x*RT for NP1, 3.45*±*0.08 *x*RT for NP2; `WavPack`: 12.82*±*0.57 *x*RT for NP1, 12.27*±*0.45 *x*RT for NP2). Compared to the `Zarr`-based codecs, one can then expect around an 8% reduction in file size when using audio compression for both probe types (NP1 - `blosc-zstd`: *∼*35.8%, `LZMA`: *∼*35.4%; NP2 - `blosc-zstd`: *∼*52.63% 1.9*±*0.06, `LZMA`: *∼*52.4%). For applications in which decompression speed is critical—for example if the data needs to be accessed repeatedly with low latency—`blosc-zstd` appears to provide a good compromise between compression ratio and decompression speed (note that in Figure 6c and 6f, the *y*-axis is plotted on a logarithmic scale).

**Figure 6:**
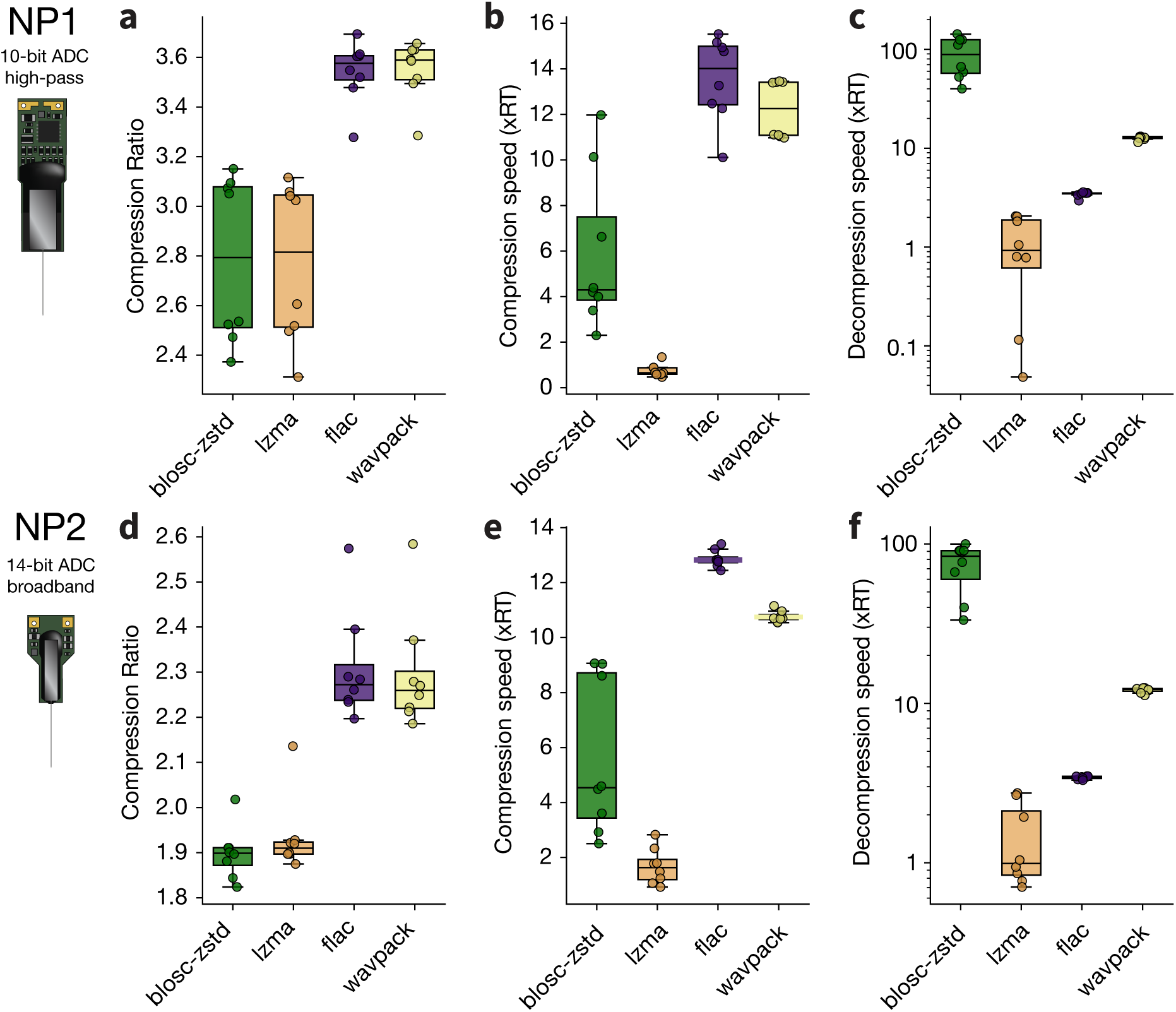
Comparing audio codecs and the highest performing general-purpose codecs. Compression metrics for two general-purpose codecs, `blosc-zstd` (green) and `LZMA` (orange), and the two audio codecs, `FLAC` (purple) and `WavPack` (yellow). Reported values are based on the *high* compression level for `blosc-zstd` and `LZMA`, and the *medium* compression level for `FLAC` and `WavPack`. Bit shuffling is applied for `blosc-zstd`, while for `LZMA` we report the shuffling options that yielded best compression rations for the two probes: no shuffling for NP1 and bit shuffling for NP2. The top row (**a-c**) is for NP1, the bottom row (**d-f**) is for NP2. *N* = 8 compression jobs for all distributions; individual data points are shown.

Finally, we looked at the effects of delta filters and pre-processing options (Figure 7). While delta filters do not improve compression for audio codecs (`WavPack` compression is even impaired with *1d* delta filtering), filtering in the time dimension (*2dT*) appears to improve compression ratios for general-purpose codecs (the same is true, to a lesser extent, for delta filters in time and space). With a time delta filter, compression ratio for `blosc-zstd` increases from 2.79*±*0.33 to 3.24*±*0.09 for NP1 and from 1.9*±*0.06 to 2.13*±*0.12 for NP2, reducing the gap with the audio codecs, which still provide the highest compression ratios. Nevertheless, the additional processing required to reconstruct the signals from their delta representation at decompression strongly impacts decompression speeds, which for *2dT* is reduced to 8.03*±*0.15 *x*RT and 8.03*±*0.23 *x*RT for NP1 and NP2, respectively (Figure S4), making decompression with this option slower than using `WavPack` on its own.

**Figure 7:**
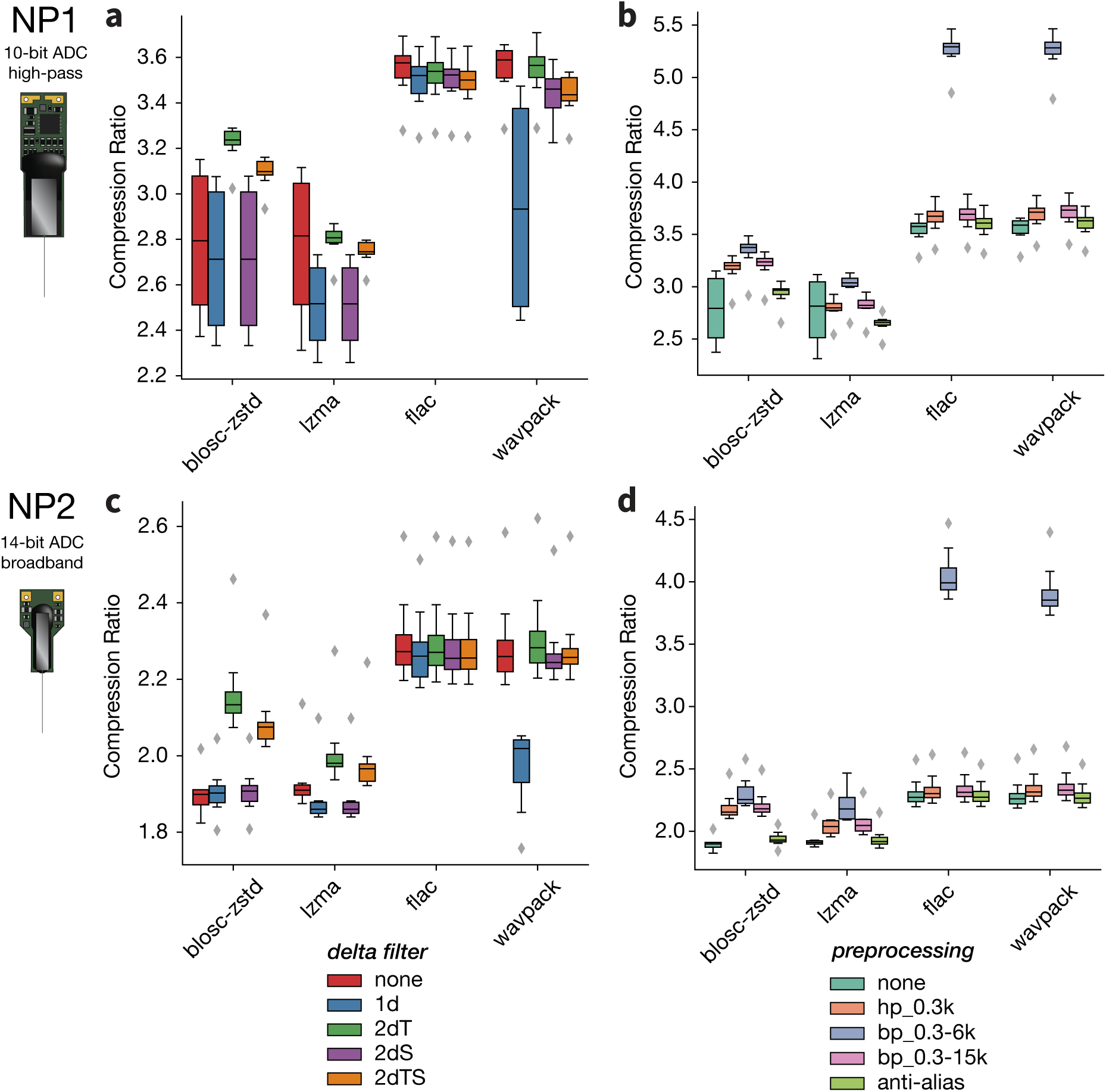
Effect of delta filters and pre-processing options. Compression ratios for `blosc-zstd`, `LZMA`, `FLAC`, and `WavPack` after applying different delta filters (**a** and **c**) or pre-processing steps (**b** and **d**). The top row (**a** and **b**) is for NP1, the bottom row (**c** and **d**) is for NP2. *N* = 8 compression jobs for all distributions.

Applying a high-pass filter slightly improves compression performance for all codecs and probes (Figure 7b,d). This is also true for NP1, even though the *AP* stream includes an analog 300-Hz high-pass filter. This is probably due to the higher order of the digital filter applied here, which attenuates low frequencies with more efficacy. Although NP2 data is broadband, the application of a high-pass filter prior to compression only marginally increases compression ratio, which may indicate that the higher ADC resolution, and not the lack of an analog high-pass filter, is the main reason for the lower compression ratios achieved for NP2 data. The application of the 0.3-6 kHz band-pass filter, especially for audio codecs, greatly increases compression ratio in all cases, suggesting that high-frequency components of electrophysiology signals are harder to efficiently compress than low frequencies. For `WavPack`, for example, the compression ratio goes from 3.59*±*0.12 for NP1 and 2.27*±*0.13 for NP2 when no pre-precessing is applied, to 5.28*±*0.2 for NP1 and 3.85*±*0.22 for NP2 after band-pass filtering. The application of a band-pass filter with a low-pass cutoff at the Nyquist frequency (15 kHz) only marginally increases compression ratio with respect to a high-pass filter with a cutoff at 300 Hz. Applying an anti-aliasing filter between 0.5 Hz and 15 kHz results in an even lower or null compression ratio increase: for `WavPack`, for example, the compression ratio increases to 3.61*±*0.13 for NP1, but no benefit is observed for NP2 (2.26*±*0.11).

### Lossy compression

After comparing the available lossless compressors, we investigated *lossy* compression using two strategies: bit truncation (using `blosc-zstd` as the base codec) and `WavPack` Hybrid mode (see Lossy compression in the Methods section). We used both experimental and simulated datasets for this comparison. To limit the necessary computation time, we restricted the analysis to four experimental datasets for each probe type.^16^

All benchmarks, which included spike sorting with `Kilosort2.5`, were run on a 16-CPU machine with 64 GB of RAM and a GPU (AWS `g4dn.4xlarge` EC2 instance) through the Code Ocean computing platform. Compression jobs were run using 16 parallel workers.

#### How much can we gain from lossy compression?

We first look at how much, in terms of compression performance, we can gain if we employ a lossy compression strategy (Figure 8). For bit truncation, compression ratio distributions for NP1 and NP2 do not overlap, due to the higher ADC bit depth of NP2 compared to NP1 (14 bits vs. 10 bits). For NP1, right-shifting each sample by 3-4 bits results in compression ratios between 5 and 10, while for larger truncations the compression ratio dramatically increases, which likely indicates that most of the samples are cut to zeros (Figure 8a). This is also reflected in the root-mean-square error (RMSE) between the original and truncated samples, which increases to values beyond 15 µV for bit truncations above 4 (Figure 8c). Considering that the nominal value of input-referred noise for NP1 is reported to be 5.9 µV, such large values indicate that most of the signal is lost. For NP2, given its higher ADC depth and dynamic range, the RMSE curve appears to be shifted to the right by 2 bits relative to that of NP1 (Figure 8c). This is due to the fact that NP2 signals have a resolution 4 times higher than NP1 *AP* streams. However, the compression ratio curve for NP2 is lower than a right-shifted version of the NP1 curve (see NP1 bit 5 versus NP2 bit 7 in Figure 8a). This discrepancy is likely due to the broadband nature of the NP2 signals.

**Figure 8:**
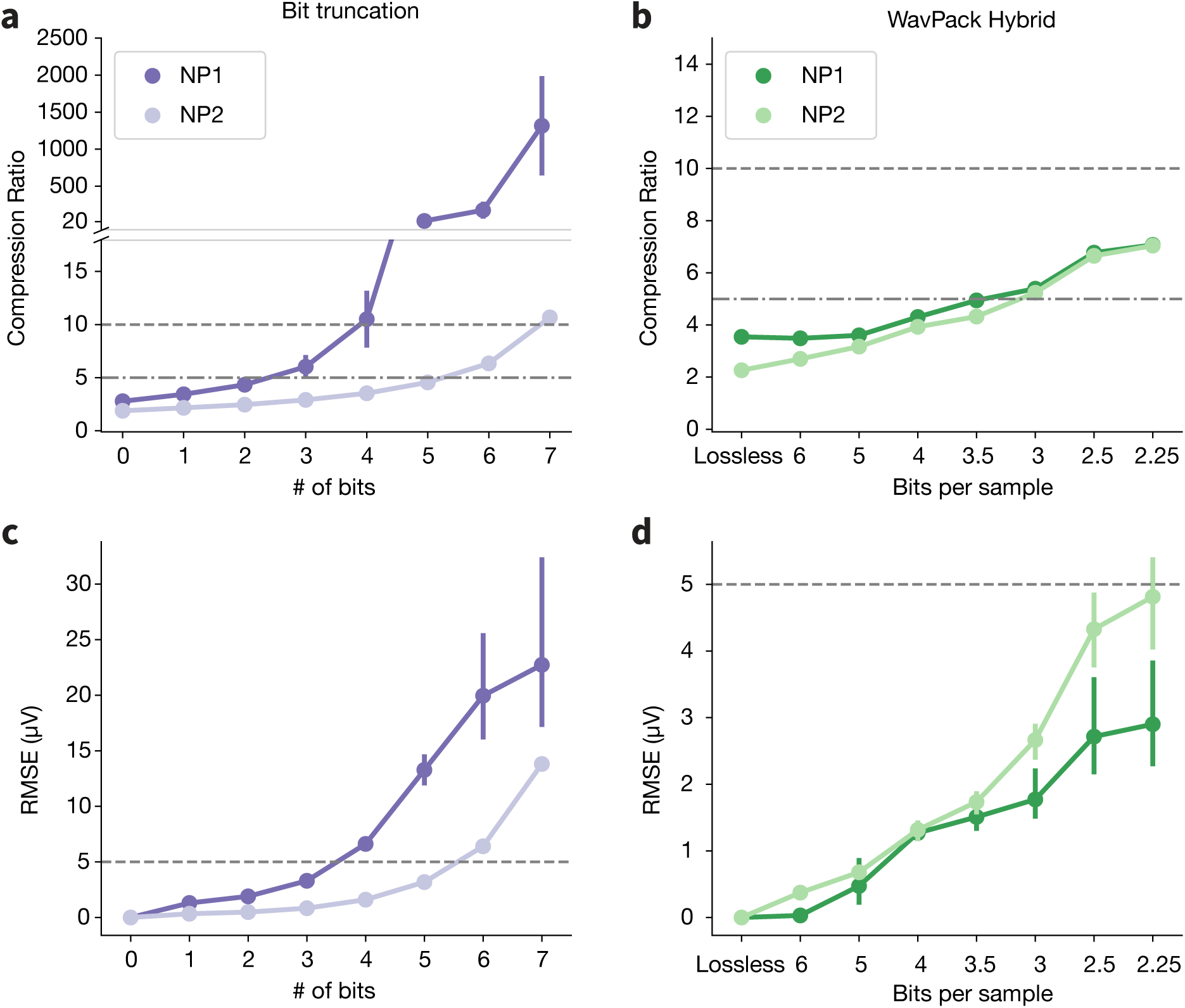
Impact of lossy compression on experimental data. Compression ratios (**a-b**) and root-mean-square error (RMSE) values (**c-d**) for bit truncation (left, purples) and `WavPack` Hybrid (right, greens). The *x*-axes represent the number of bits truncated (for bit truncation) or the target *bps* (for `WavPack` Hybrid). In all panels, lossiness increases from left to right. Horizontal dashed lines are added for reference and mark compression ratios of 5 and 10 or an RMSE of 5 µV. *N* = 4 compression jobs for each condition.

The `WavPack` Hybrid approach (Figure 8b,d) shows similar compression ratios for NP1 and NP2 data, with curves for both probes reaching a compression ratio slightly above 7 (file size *∼*14%) for a *bps* value of 2.25 (NP1: 7.08*±*0.02, NP2: 7.04*±*0.01). The RMSE curves remain below 5 µV for all values. The NP2 errors are slightly higher than the NP1 ones, again probably due to the larger dynamic range of NP2 signals. Note that for NP1 and *bps ≥* 5, the Hybrid compression is effectively lossless (Figure 8b). This can be explained by the fact that `WavPack` can already compress losslessly with a CR of 3.59, i.e., the 16-bit samples can be already encoded, on average, at 16/3.59*≈*4.45 bits per sample.

The highest `WavPack` Hybrid compression ratio, therefore, is similar to that achieved by truncating between 3 and 4 bits for NP1 and between 6 and 7 bits for NP2 (NP1 - 3 bits: 6.07*±*1.51, 4 bits: 10.64*±*17.71; NP2 - 5 bits: 4.64*±*1.71, 6 bits: 11.2*±*0.87). Similar trends of compression ratios were obtained using simulated data (Figure S5a-b). In simulations, however, the NP2 data only contain the *AP* band, which explains the different values of compression ratio for high bit truncation values (6-7) compared to experimental data (see Figure 8a). The `WavPack` Hybrid behavior on simulated data, instead, appears to match almost perfectly that obtained for experimental data. Finally, RMSE values are lower for simulated data, which can be explained by the lower RMS noise values of simulated data with respect to experimental data (Figure S5c-d).

To directly visualize the effects of lossy compression, we show 10 ms snippets of the voltage traces (after a bandpass filter and common median reference) as lossy compression becomes more aggressive (Figure 9). In Appendix 1 we provide an more detailed view of the decompressed traces generated using the `Figurl` web-based visualization plugin (Magland & Soules, n.d.).

**Figure 9:**
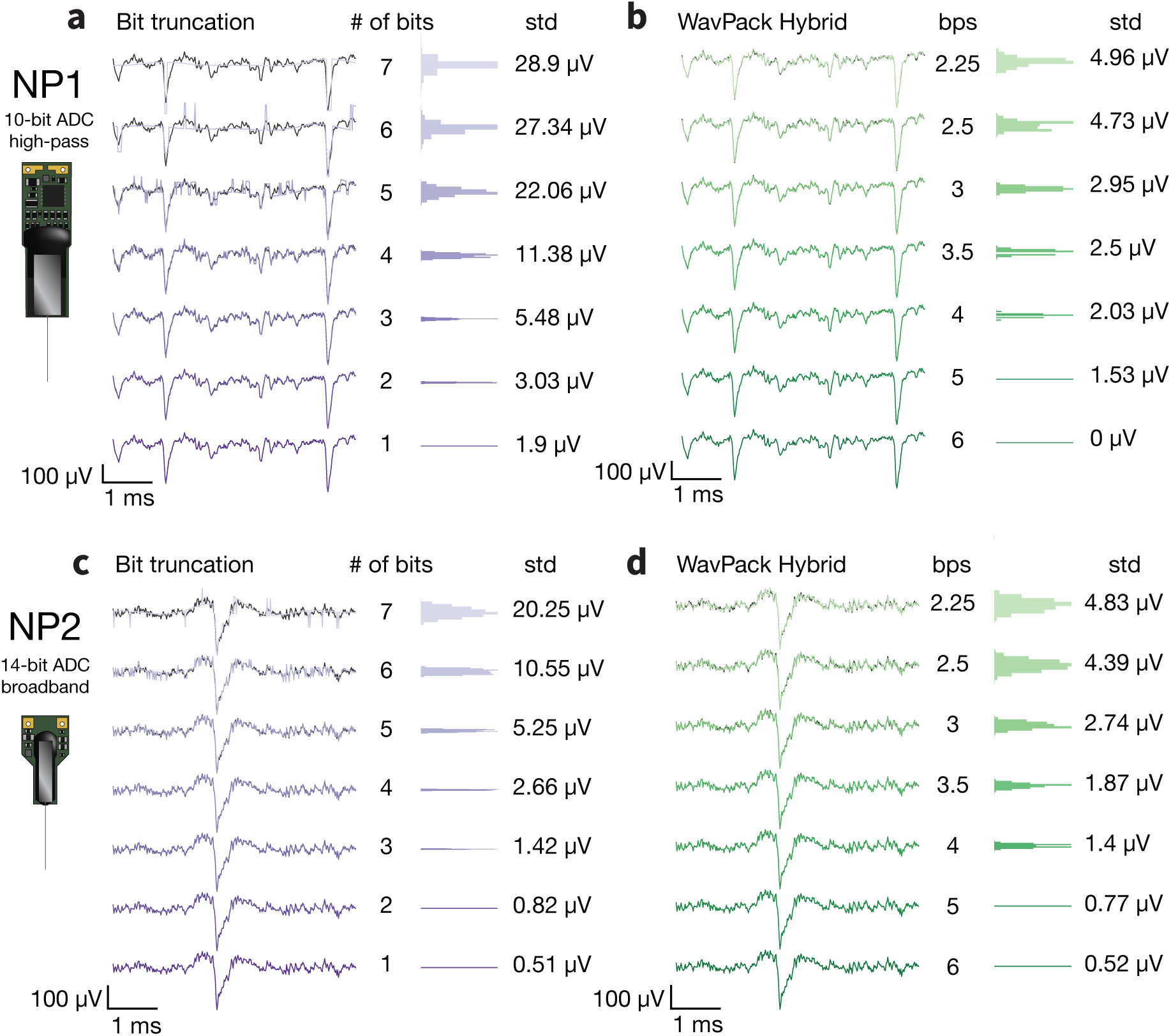
Effect of lossy compression on voltage traces. Sample 10 ms traces following decompression, filtering, and common median referencing. The top row (**a-b**) is an NP1 recording (625749_2022-08-03_15-15-06_ProbeA); the bottom row (**c-d**) is an NP2 recording (595262_2022-02-21_15-18-07_ProbeA). The right portion of each panel shows the distribution and standard deviation of the error of each decompressed sample relative to the original trace. Note that the scale of the error distributions differs for bit truncation and `WavPack` Hybrid. In all panels, lossiness decreases from top to bottom.

#### Effects of lossy approaches on spike sorting

If it is to be used in practice, lossy compression must not degrade spike sorting results. To check whether lossy compression impacts spike sorting, we first used simulated data with known spiking activity. For these datasets, we can compute accuracy, precision, and recall metrics for each ground truth spike train (Figure 10). These three performance metrics appear to be unaffected for all *bps* values and both probe types when using the `WavPack` Hybrid strategy. For bit truncation, spike sorting NP1 data seems slightly affected after 5 bits are truncated and is drastically impaired for 6 and 7 bits. For NP2 probes, 7 bits must be truncated before spike sorting performance is reduced, consistent with the 4*×* higher voltage resolution of these devices.

**Figure 10:**
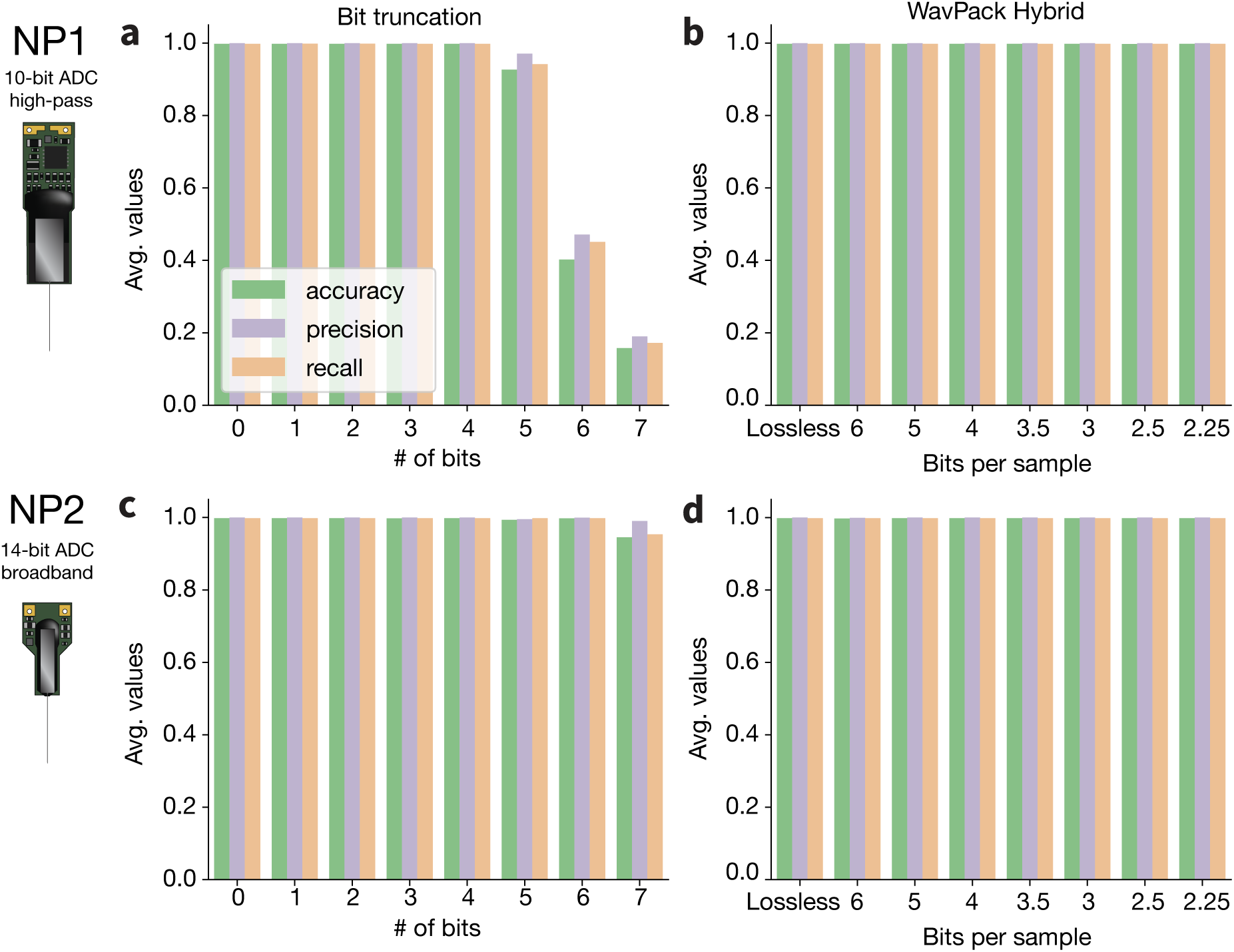
Spike sorting performance following lossy compression of simulated data. Average accuracy, precision, and recall for sorted units after bit truncation (**a** and **c**) or compression with `WavPack` Hybrid (**b** and **d**). Top row is for NP1, bottom row is for NP2.

Accuracy, precision, and recall reflect the ability of a spike sorter to correctly detect ground truth units, but they do not provide information about incorrectly identified units. To provide a more complete overview of spike sorting performance, we analyzed the number of well-detected, false positive, redundant, and overmerged units (Figure 11; see definitions in the Evaluation and benchmarks section). For example, for 5 bits truncated for NP1, the spike sorting metrics are only slightly affected (Figure 10a), but for the same condition, the number of false positive and redundant units explode (Figure 11a). The exact same situation is also observed for 7 truncated bits for NP2 (Figure 11c). For the `WavPack` Hybrid approach, the results confirm that spike sorting is not adversely affected, with a relatively constant number of well-detected and redundant units for all *bps* values (Figure 11b,d). Decreasing the target bit rate does seem to reduce the number of false positive units for NP2, presumably because compression has a slight denoising effect.

**Figure 11:**
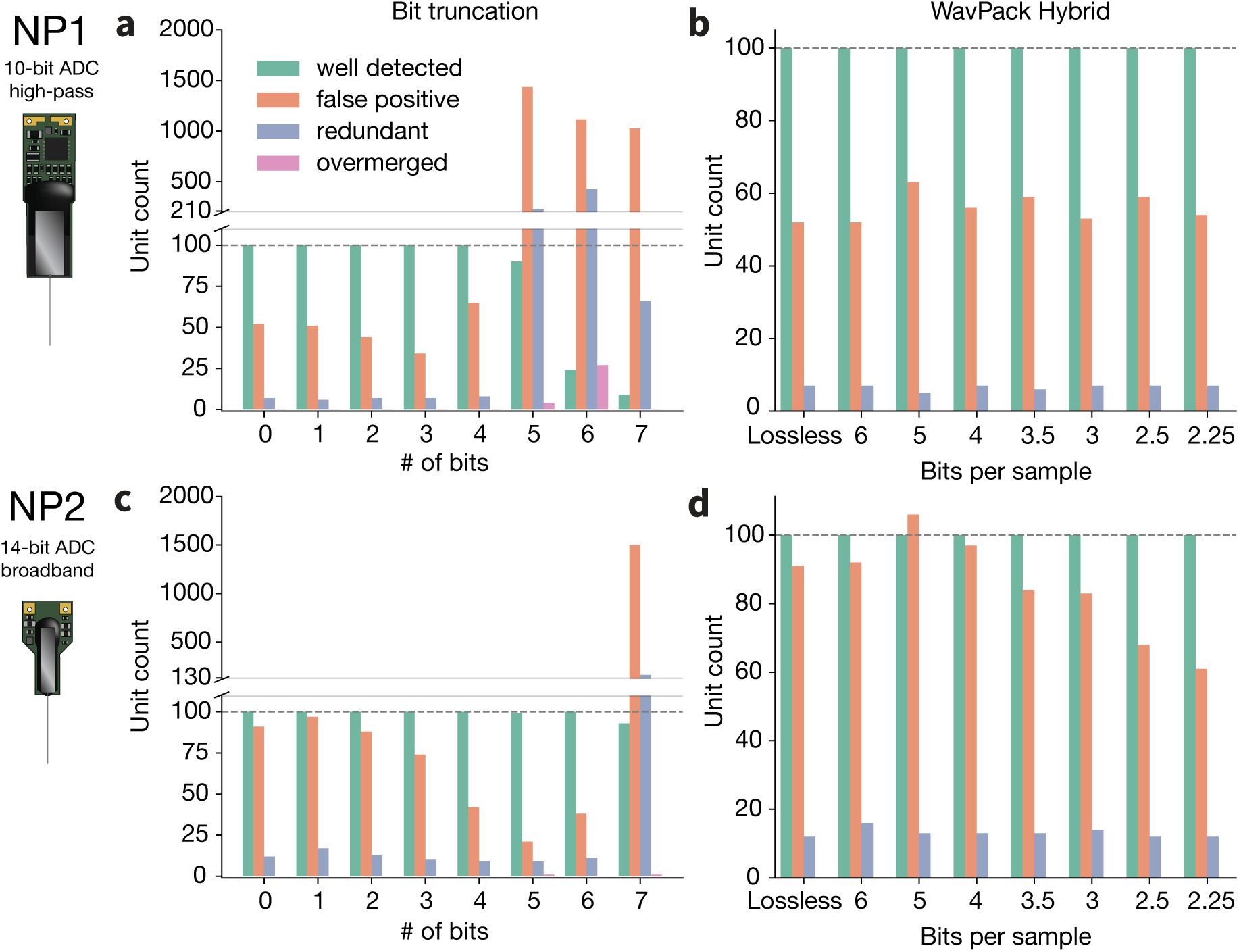
Classification of units detected following lossy compression of simulated data. Number of well detected, false positive, redundant, and overmerged units after bit truncation (**a** and **c**) or compression with `WavPack` Hybrid (**b** and **d**). Top row is for NP1, bottom row is for NP2.

The results from our simulations suggest that the `WavPack` Hybrid approach does not affect spike sorting outcomes. To validate this on experimental data, where ground truth spike trains are not available, we automatically curated the spike sorting outputs of eight sessions (see the Evaluation and benchmarks section) and counted the fraction of *passing* and *failing* units compared to the lossless spike sorting result (Figure 12). We observed similar trends as in simulated data. For NP1, the number of failing units increases dramatically for more than 5 bits truncated, and the number of passing units is reduced after 4 bits. For NP2, the number of passing and failing units both decrease for 6 and, more pronouncedly, 7 truncated bits. The fractions of passing and failing units, instead, appear generally constant for `WavPack` Hybrid, with a slight reduction in the number of passing and failing units for the lowest *bps* values (2.5 and 2.25).

**Figure 12:**
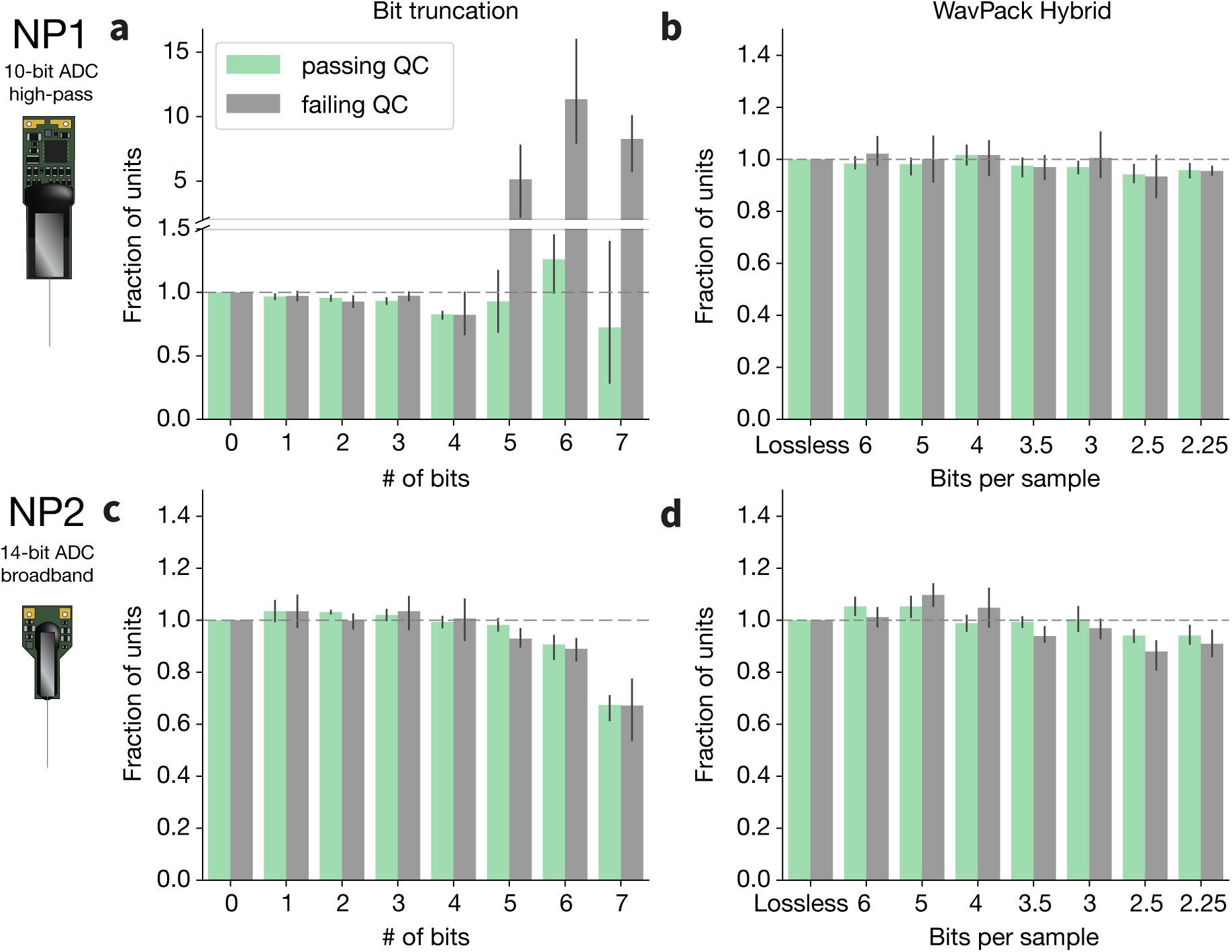
Spike sorting performance following lossy compression of experimental data. Fraction of units passing or failing quality control compared to the spike sorting results on uncompressed data after bit truncation (**a** and **c**) or compression with `WavPack` Hybrid (**b** and **d**). Top row is for NP1, bottom row is for NP2. *N* = 4 spike sorting runs for each condition.

To further investigate how spike sorting results are affected by lossy compression, we compared the full spike trains for all units passing QC against the best matching units detected in the lossless data. In Figure 13, each line represents the fraction of spike times in agreement between different lossy sortings and the lossless scenario. Even for two separate spike sorting runs applied to the same lossless data (the “0” line for bit truncation and the “Lossless” line for `WavPack` Hybrid), the detected spike trains do not match perfectly, as indicated by the sloping of the agreement lines toward zero (an exact match would result in a flat line at 1). This is attributed to the inherent run-by-run variability for `Kilsort 2.5`. Increasing the number of truncated bits quickly reduces the agreement with respect to the lossless result for NP1 (Figure 13a), while the agreements for NP2 only seem to be affected only after 6 and 7 truncated bits (Figure 13c). The abrupt drops in the agreements are due to partial matches and over-splits. For `WavPack` Hybrid, spike sorting results are more stable (Figure 13b,d), but some divergence from the lossless output appears for *bps* of 2.5 and 2.25. However, for other datasets, the divergence can arise at higher *bps* values (Figures S6 and S7). The same divergence can be also observed when comparing all detected units, rather than only those passing QC (Figure S8a,c).

**Figure 13:**
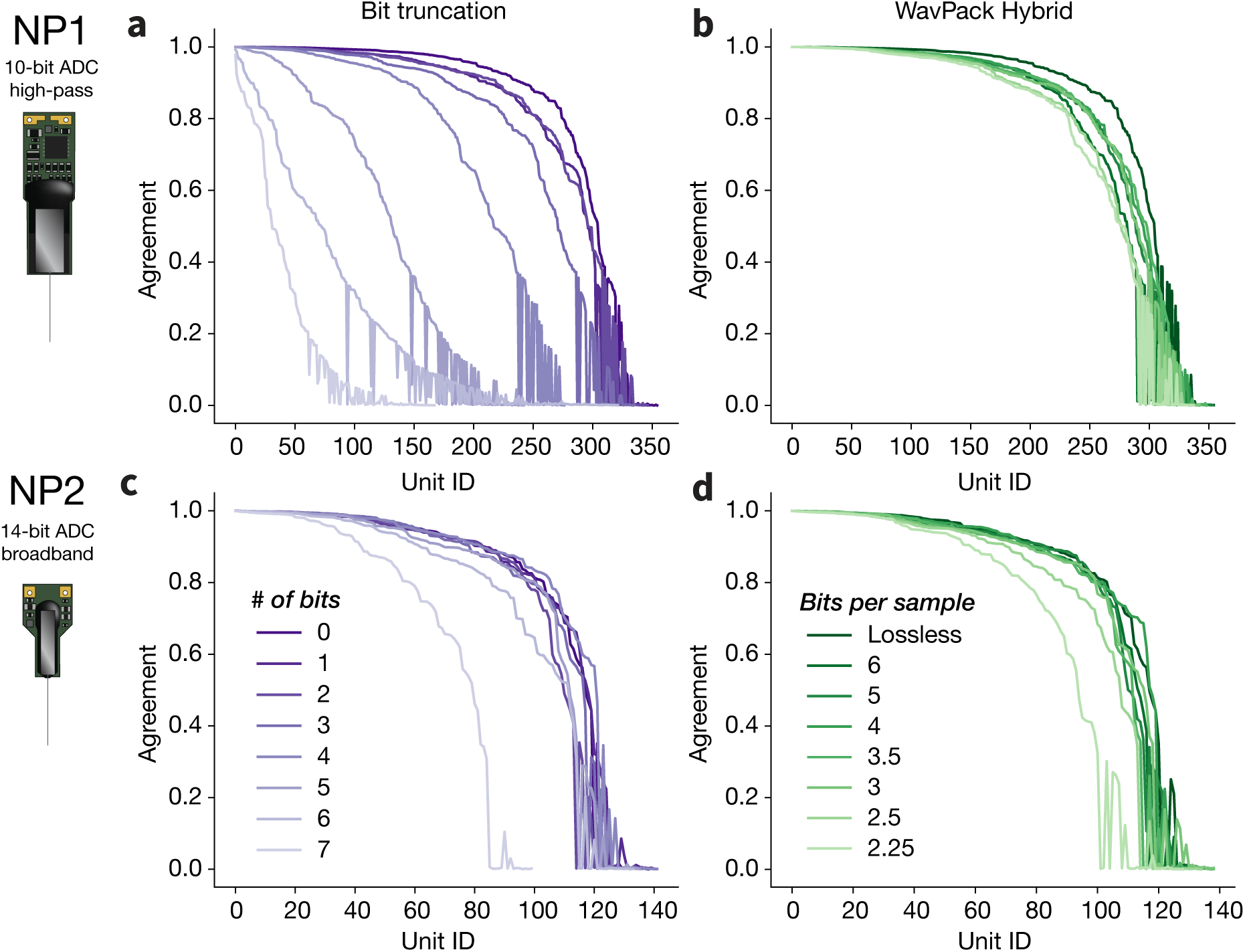
Comparing spike trains detected before and after lossy compression. Each line represents the ordered agreement of matching units between the lossy and lossless spike sorting outputs. If all spike times are identical, units will have an agreement of 1. For the 0 bits truncated or lossless `WavPack` compression, we compared the outputs of two `Kilosort 2.5` runs on the same data. The top and bottom rows display NP1 (CSHZAD026_2020-09-04_probe00) and NP2 (612962_2022-04-13_19-18-04_ProbeB) results, respectively. Left panels (**a** and **c**, purples) represent bit truncation and right panels (**b** and **d**, greens) are for `WavPack` Hybrid compression.

We investigated whether these discrepancies might arise because the spike sorter consistently finds a larger or smaller number of spikes after compression. To do so, we first compared the spike trains from the lossless compression with the ones from `WavPack` Hybrid with a *bps* of 2.25. Next, we selected all matching unit pairs with an agreement above 0.5 and, for each of these pairs, we counted the fraction of *excess* spikes in the lossy or lossless spike trains. The distributions of the fraction of excess spikes are symmetric (Figure S8b,d), which confirms that lossy compression does not cause the spike sorter to consistently miss spikes, nor consistently find additional spikes with respect to the lossless sort. Sample waveforms of excess spikes from a matched unit pair are visually indistinguishable (Figure S8b,d, inset), suggesting that the observed differences in spike sorting results are likely to be attributed to inherent run-to-run variability, rather than to systematic modulation of waveforms due to lossy compression.

#### Waveform preservation for lossy compression strategies

Preserving spike waveforms is critical for accurate spike sorting, and waveform features carries useful information about neurons’ underlying biophysical properties. To further assess whether lossy compression distorts spike waveforms, we computed the relative errors of three commonly used features (peak-to-valley duration, half-width duration, and peak-to-trough ratio) with respect to the ground truth average waveforms in our simulated datasets. For each unit (*N* = 100), these features were computed on the *main* channel (the one with the largest amplitude) and on a *peripheral* electrode 60 µm away (Figure 14). Truncating 3 to 4 bits for NP1 and 6 to 7 bits for NP2 results in errors well above 20%, especially for peripheral channels. The `WavPack` Hybrid strategy, on the other hand, preserves waveform features with much higher fidelity, with all error distributions remaining well below 10%, and staying largely constant with increasing *bps*. In terms of compression ratio, `WavPack` Hybrid with a *bps* of 2.25 is equivalent to truncating between 3 and 4 bits for NP1 and 6 and 7 bits for NP2. Nevertheless, lossy compression with `WavPack` Hybrid has a minimal impact on waveforms.

**Figure 14:**
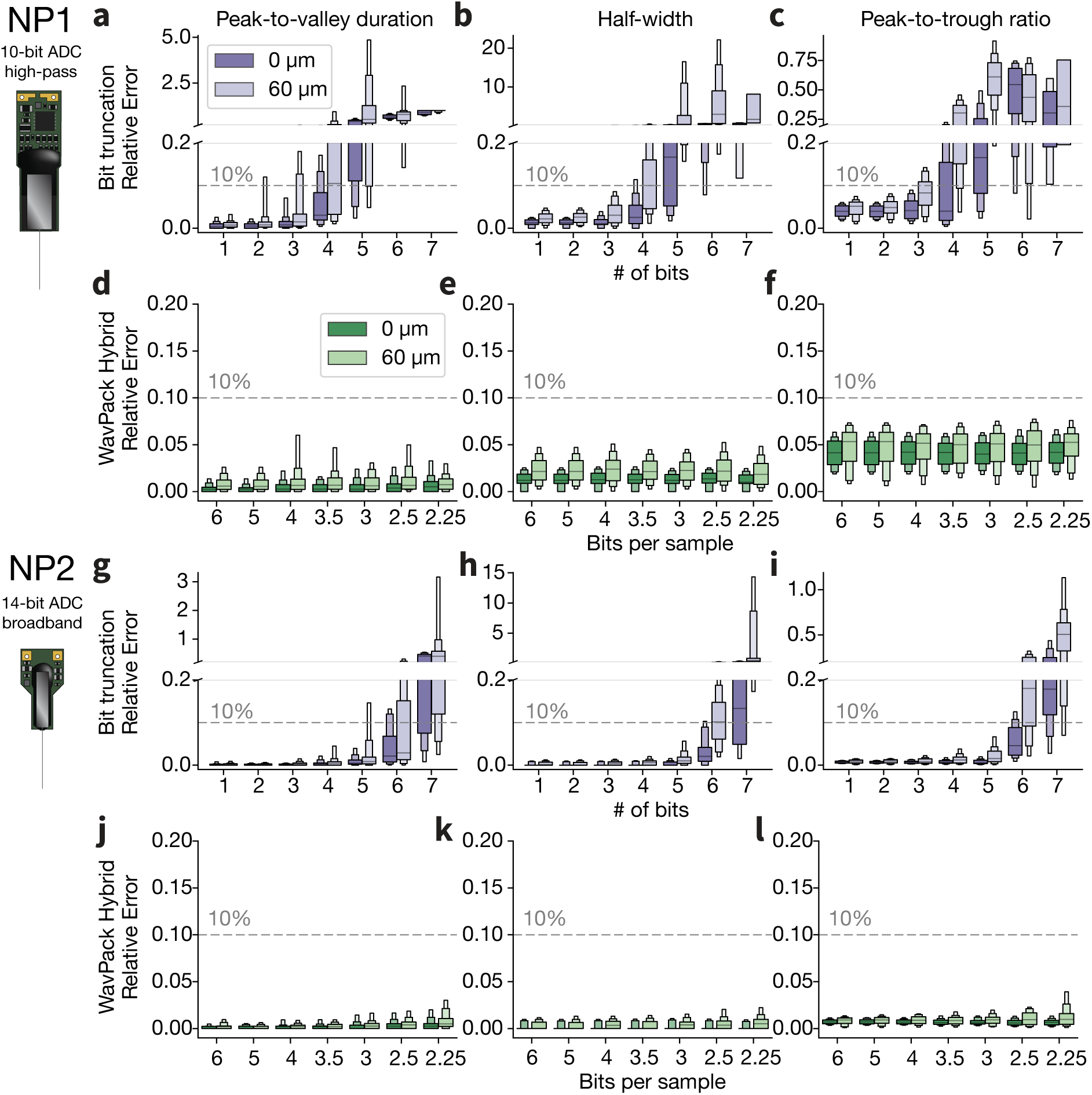
Effect of lossy compression on spike waveforms. Relative errors in estimating common waveform features (from left to right columns: peak-to-valley duration, half-width duration, and peak-to-trough ratio) for NP1 (**a-f**) and NP2 (**g-l**) on simulated data. For each metric, values are computed on the main channel (darker hues, marked as 0 µm) and on a peripheral channel at 60 µm distance from the main channel (lighter hues). For reference, the horizontal dashed lines represent a 10% relative error. *N* = 100 units for all error distributions.

### Software extensions

As part of this project, we developed and shared a range of software tools. The following sections summarize our contributions in this area.

### Integration of Zarr-based compression in SpikeInterface

To facilitate the compression of electrophysiology data, we extended the `SpikeInterface` framework (Buccino et al. 2020) to support a `Zarr` backend (SI version 0.94 and higher). `SpikeInterface` natively reads over 30 common data acquisition formats (including `Open Ephys` and `SpikeGLX`), which can now be easily loaded and written to disk as compressed `Zarr` files. We used this `Zarr` backend to systematically evaluate and benchmark all the compression strategies presented in the Lossless compression and Lossy compression sections.

This code snippet shows how to compress an Open Ephys dataset with the spikeinterface.save() function, using parallel processing (8 jobs) and chunked writing (1 s chunk size):

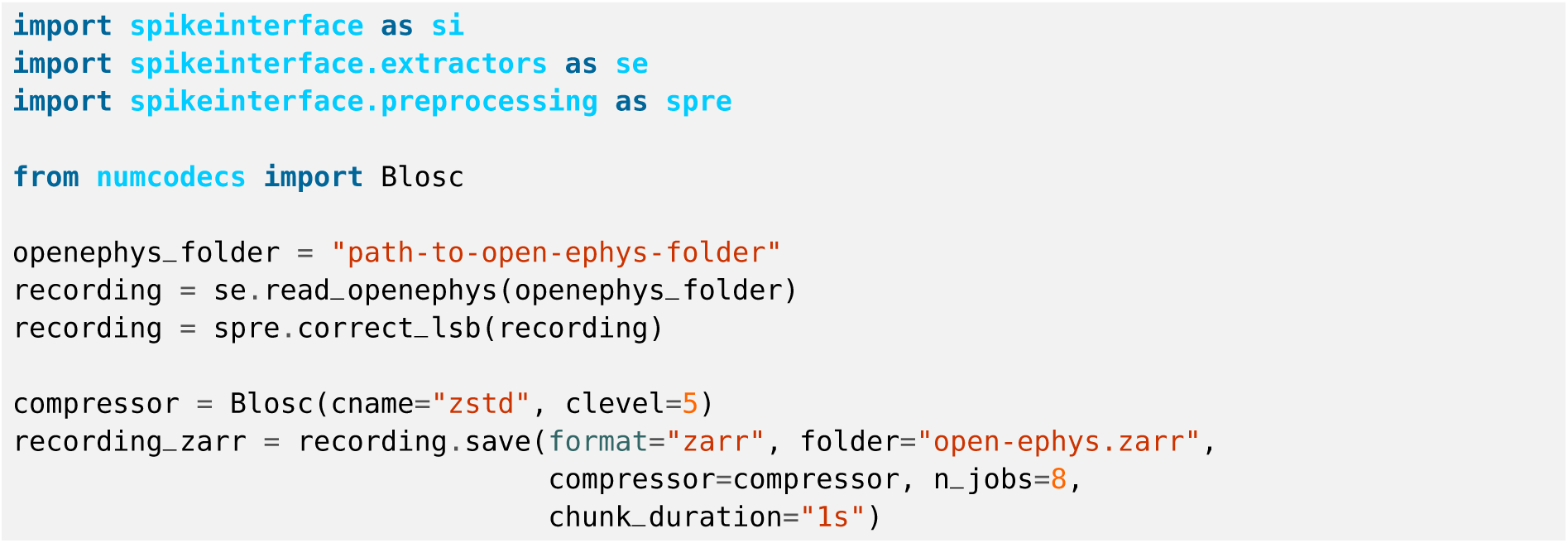

In this example, LSB correction (which we assessed in the Lossless compression section) is enabled via the correct^_^lsb() function.

Because Zarr supports writing directly to cloud storage, the same interface can be used to save a local dataset directly to a cloud bucket, just by specifying the appropriate access options:

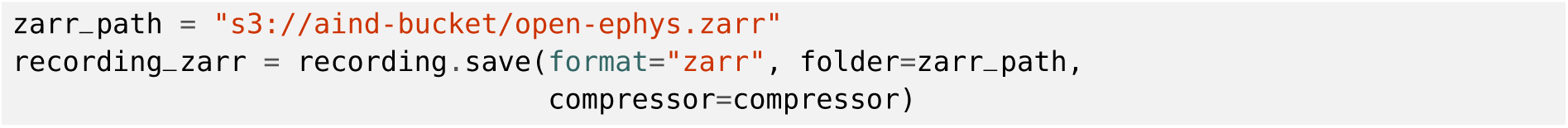

We assume here that the local machine has the correct access credentials to write to the s3://aind-bucket storage location.

`Zarr` datasets saved by `SpikeInterface` and stored on the cloud can be accessed **lazily**, i.e., only the requested chunks are downloaded and decompressed:

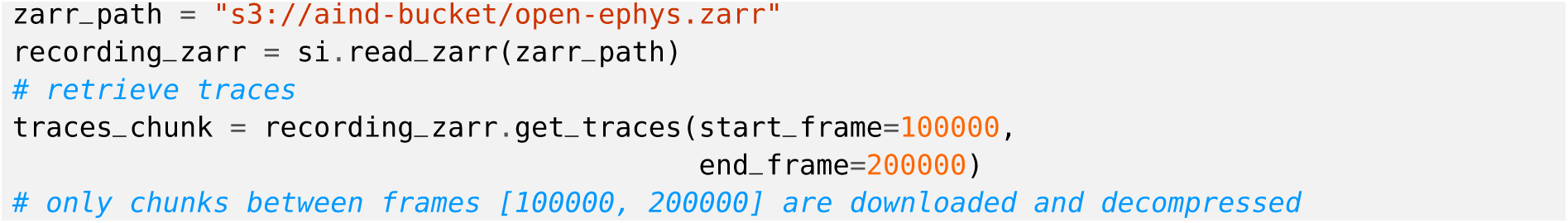

This facilitates efficient access to the data within a cloud computing environment.

### Extensions to the numcodecs library

As part of this project, we developed three `numcodecs` extensions: the `FLAC` and `WavPack` *audio* codecs and a `Delta2D` filter. The new codecs are open source, with the source code available on GitHub (`flac-numcodecs`, `wavpack-numcodecs`, and `delta2D-numcodecs`) and `Python` packages on `PyPi`. The three packages are cross-platform (tests are run on Windows, MacOS, and Ubuntu with Continuous Integration) and can be installed directly with `pip`:

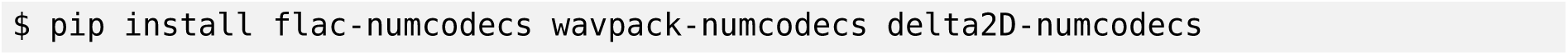

Once installed, the *audio* compressors are automatically added to the `numcodecs` registry and can be used directly in `Zarr` or `SpikeInterface`. In this example, we show how to compress a recording with `FLAC` (compression level = 8 – *high*, channel block size = 2), `WavPack` lossless mode (compression level = 3 – *high*), and `WavPack` hybrid mode (*bps* = 3), as well as with a delta filter in the time dimension using the `blosc-zstd` compressor:

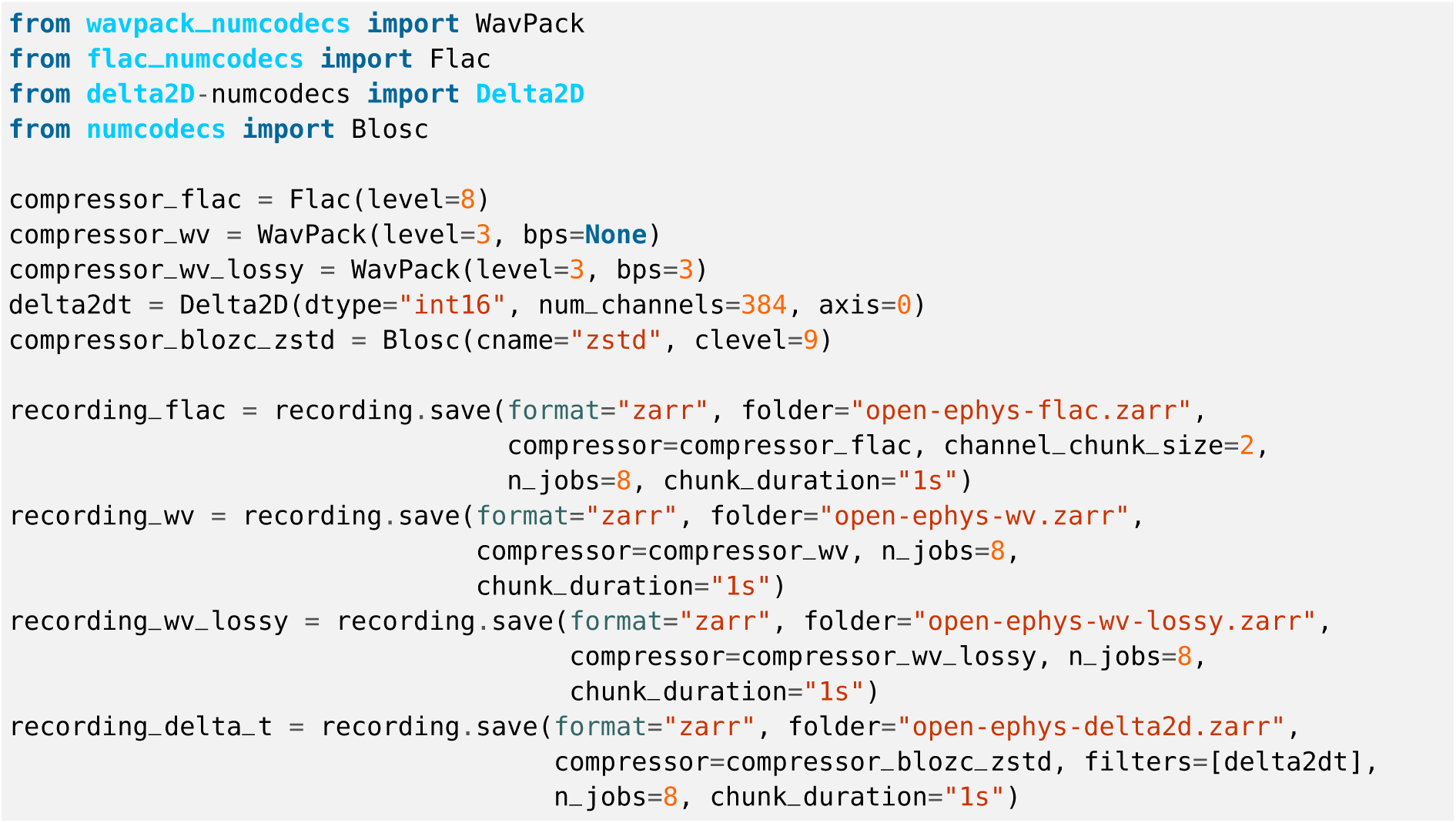

### Integration with NWB

All the compression strategies that we utilized (and developed) in this contribution can be used to compress electrophysiology data in the Neurodata Without Borders (NWB) format (Teeters et al. 2015; Rübel et al. 2022) through the recently developed `Zarr` backend (Tritt et al. 2019).^17^ The `FLAC` codec, but not `WavPack`, is also available as an `HDF5` Registered Filter Plugin.^18^

## Discussion

We have presented a detailed comparison of compression strategies for large-scale electrophysiology data. In addition, we implemented a `Zarr`-based compression framework within the `SpikeInterface` API, which is now available for the community to use. Below, we consider the implications of our findings and provide recommendations for other researchers collecting similar types of data.

### Lossless compression

We compared 11 different lossless codecs across multiple types of recordings and compression options (including chunk size, compression level, and pre-shuffling). We found that audio codecs (`FLAC` and `WavPack`) outperform general-purpose codecs in terms of compression ratio (CR), allowing one to compress Neuropixels 1.0 data up to a CR of 3.59*±*0.12 (file size *∼*28%), and Neuropixels 2.0 data up to a CR of 2.27*±*0.13 (file size *∼*44%). The downside of audio compressors is that they have relatively slow decompression speeds (e.g., *∼*12 *x*RT for `WavPack` vs. *∼*50-90 *x*RT for `blosc-zstd`). However, the reported decompression speed is computed by decompressing 10 s of traces without explicit parallelization. In most cases, decompression (i.e., reading the data) only needs to be performed a handful of times and can utilize multiple CPU cores. If the parallel reading (and processing) of the compressed data matches the chunk size (e.g., 1 s), then one can expect a speed-up proportional to the number of parallel workers. The very high speed of the natively multi-threaded `blosc-zstd` codec (Figure 6c,f) could make this the best option for applications that require frequent decompression, such as trace visualization. When using general-purpose compressors, applying an additional delta filter in the time dimension can further boost compression performance (Figure 7a-b), but would also reduce decompression speed to less than 10 *x*RT (Figure S4a-b).

In the Evaluation and benchmarks section, we argued that compression ratio and decompression speed are the main metrics to consider when evaluating compression performance. Compression ratio is inversely proportional to the price of long-term data storage, but decompression requires compute resources, which have a nonzero cost. How much, then, can we actually expect to save by using `WavPack`, which has a higher compression ratio, instead of `blosc-zstd`, which can decompress faster? To take a concrete example, we compare the costs associated with lossless compression of one hour of NP1 data (80 GB). Assuming storage at $0.023/GB per year, `blosc-zstd` will produce a compressed file of 28.67 GB (average CR = 2.79) that will cost $7.91/year to store. `WavPack` will compress the same data down to a 22.28 GB file (average CR = 3.59) which will cost $6.15/year. Assuming a cost of $0.05/CPU per hour,^19^ and assuming decompression occurs at the median speeds reported in Figure 6c (88.89 *x*RT for `blosc-zstd` and 12.82 *x*RT for `WavPack`), `blosc-zstd` would become cheaper overall only if the full dataset were decompressed at least 528 times in a year (with similar calculations, the number of required decompressions for NP2 would be 532).^20^ Since we expect a typical cloud analysis pipeline will decompress the data only prior to pre-processing and spike sorting and to generate, at that time, appropriate visualizations for quality control and inspection, we conclude that `WavPack` is the more cost-effective choice. While the raw data may need re-processing, for example when testing new methods for pre-processing or spike sorting, we expect this to happen, at most, a few times a year, well below the 500+ decompressions needed to reach the break-even point.

We also showed that pre-processing with a high-pass or band-pass filter can increase compression ratio (Figure 7c-d). Although we included the effect of pre-processing in the lossless section, it’s important to note that pre-processing effectively results in lossy compression, since the frequencies that are filtered out cannot be recovered. While it may be tempting to apply such filters prior to compression, rather than after decompression, it is worth noting that the frequency content of spike waveforms can extend below 1000 Hz, as well as above the 6 kHz cutoff frequency used in typical band-pass filters for electrophysiology data. Several spike sorters, including `Kilosort`, pre-process the data with a high-pass filter to retain signals up to Nyquist frequencies that may help distinguish waveforms from nearby neurons. There are even attempts to develop spike sorting methods that use broadband data as input (Le Cam et al. 2023). For these reasons, we recommend compressing the raw data directly, without applying any pre-processing to boost compression ratio.

### Lossy compression

After benchmarking lossless compression, we evaluated two lossy strategies: bit truncation and `WavPack` Hybrid mode. Using simulated data, we showed that `WavPack` Hybrid is superior to naive bit truncation, as it can achieve a compression ratio of around 7 (*∼*14% file size) for both probe types without affecting spike sorting results (Figures 10 and 11) or altering spike waveforms (Figure 14). Although we observed similarly unaffected results using experimental data (Figure 12), it is harder to reach strong conclusions in this case due to the absence of ground-truth spike trains. Therefore, our findings on experimental data should be interpreted with caution, as counting the number of units that pass certain thresholds based on quality metrics does not provide a full picture of spike sorting performance. For example, even with lossless compression, spikes from a single neuron might be over-split into two units, both of which may pass quality control. We must also be wary of incomplete units, e.g. a unit that includes only 30% of the original spikes but still passes quality control.

To assess the impact of lossy compression on experimental data in more detail, we used the spike sorting results from the original (lossless) recording as “pseudo ground truth” (Figures 13, S6, and S7). In general, sorting outputs when using `WavPack` Hybrid closely matched those of the original recording, even for the highest compression levels. However, the exact *bps* value at which sorting outputs began to diverge from the original varied by experiment. While this comparison is able to reveal more subtle changes in spike sorting outputs, it may penalize lossy compression options that actually improve sorting accuracy, for example by lowering the noise floor and making spikes more easily detectable. Due to the remaining uncertainty around the precise impact of lossy compression, for our initial round of data acquisition at the Allen Institute of Neural Dynamics we are performing *lossless* compression with `WavPack`. However, given the potential for additional cost savings, it is worth pursuing further validation of lossy strategies. One option would be to assess sorting performance with hybrid recordings (Rossant et al. 2016; Wouters et al. 2021), where artificial spikes are injected into a real dataset, or paired recordings, in which the same neuron is simultaneously observed with an extracellular electrode and an intracellular pipette (Neto et al. 2016; Marques-Smith et al. 2018). An alternative approach could target downstream analysis: if the same scientific conclusions are reached from raw and lossy recordings, then one could argue that lossy compression is not affecting spike sorting in a meaningful way.

Other lossy compression strategies might also be worth exploring. Simple bit truncation, as we have utilized here, is the least computationally intensive method for reducing the bit-depth of digitally sampled signals. There are several additional compression techniques that we plan to explore in future work: *dithering*, wherein a small random signal is added to the sample values before truncation to reveal details in the signal below the least-significant bit; *noise shaping*, wherein the quantization error introduced by the truncation is high-pass filtered and included in subsequent samples; and ADPCM (Adaptive Delta Pulse Code Modulation), which is a simple scheme for representing pulse code modulated signals using a small fixed number of bits for each sample.

## Conclusion

Given the similar properties of all extracellular electrophysiology signals, we anticipate that the results we obtained with NP1 and NP2 data will generalize to different probe types and acquisition systems, as well as brain regions and species. For example, we can expect compression ratios to be inversely related to the ADC bit depth of the acquisition device and to be lower for broadband signals compared to narrower “spike-band” signals. By providing the community with a ready-to-use compression toolbox and associated benchmarks, we hope to enable electrophysiologists to employ and customize compression strategies to immediately start reducing storage costs and conserving resources for other research activities.

## Data and code availability

The raw data used in this paper and listed in Table 1 is publicly available at: s3://benchmark-data/ephys-compression through the Registry of Open Data on AWS.

The scripts used to generate the figures in this paper can be found at: https://github.com/AllenNeuralDynamics/ ephys-compression/ in the `scripts` folder. All figures can be reproduced using the following Code Ocean capsule: https://github.com/AllenNeuralDynamics/aind-capsule-ephys-compression-results, which is also available in the Code Ocean Open Science Library: https://codeocean.com/capsule/3822095/tree/v1.

## Acknowledgments

We thank the Allen Institute founder, Paul G. Allen, for his vision, encouragement, and support. We thank Anna Lakunina, Galen Lynch, Yoni Browning, and the members of the International Brain Laboratory for collecting
example datasets. We thank members of the Code Ocean team for providing support. We thank Amazon Web Services for providing cloud storage.

## Author contribution matrix

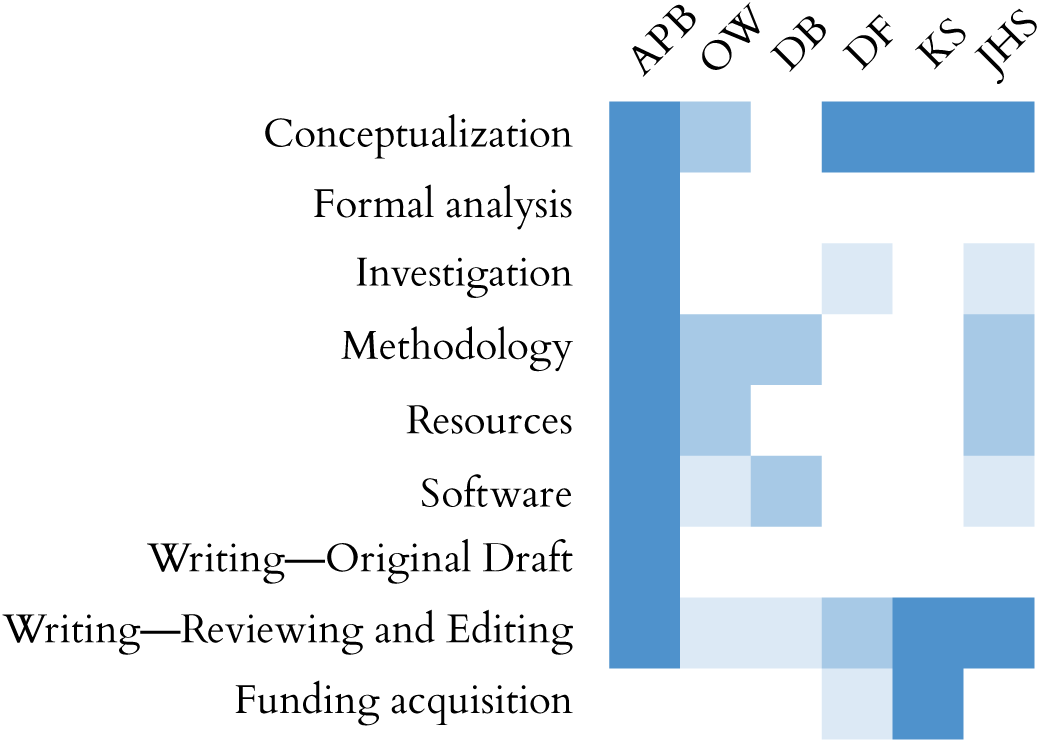

## Appendix 1. Lossy compression visualization

In this appendix, we provide Figurl-based visualizations of 1 s snippets of traces for the eight experimental datasets used for lossy compression evaluation. For each dataset and compression strategy (`WavPack` Hybrid and bit truncation), the drop-down window on the top left of the view allows one to select different layers corresponding to different levels of compression (# of bits truncated for Bit truncation, *bps* for `WavPack` Hybrid). Level 0 is the original recording in both cases.

- 595262_2022-02-21_15-18-07_ProbeA

**–** Bit truncation
**–** `WavPack` Hybrid
- 602454_2022-03-22_16-30-03_ProbeB

**–** Bit truncation
**–** `WavPack` Hybrid
- 612962_2022-04-13_19-18-04_ProbeB

**–** Bit truncation
**–** `WavPack` Hybrid
- 618384_2022-04-14_15-11-00_ProbeB

**–** Bit truncation
**–** `WavPack` Hybrid
- 625749_2022-08-03_15-15-06_ProbeA

**–** Bit truncation
**–** `WavPack` Hybrid
- 634568_2022-08-05_15-59-46_ProbeA

**–** Bit truncation
**–** `WavPack` Hybrid
- CSHZAD026_2020-09-04_probe00

**–** Bit truncation
**–** `WavPack` Hybrid
- SWC054_2020-10-05_probe00

**–** Bit truncation
**–** `WavPack` Hybrid

---

**Figure S1:**
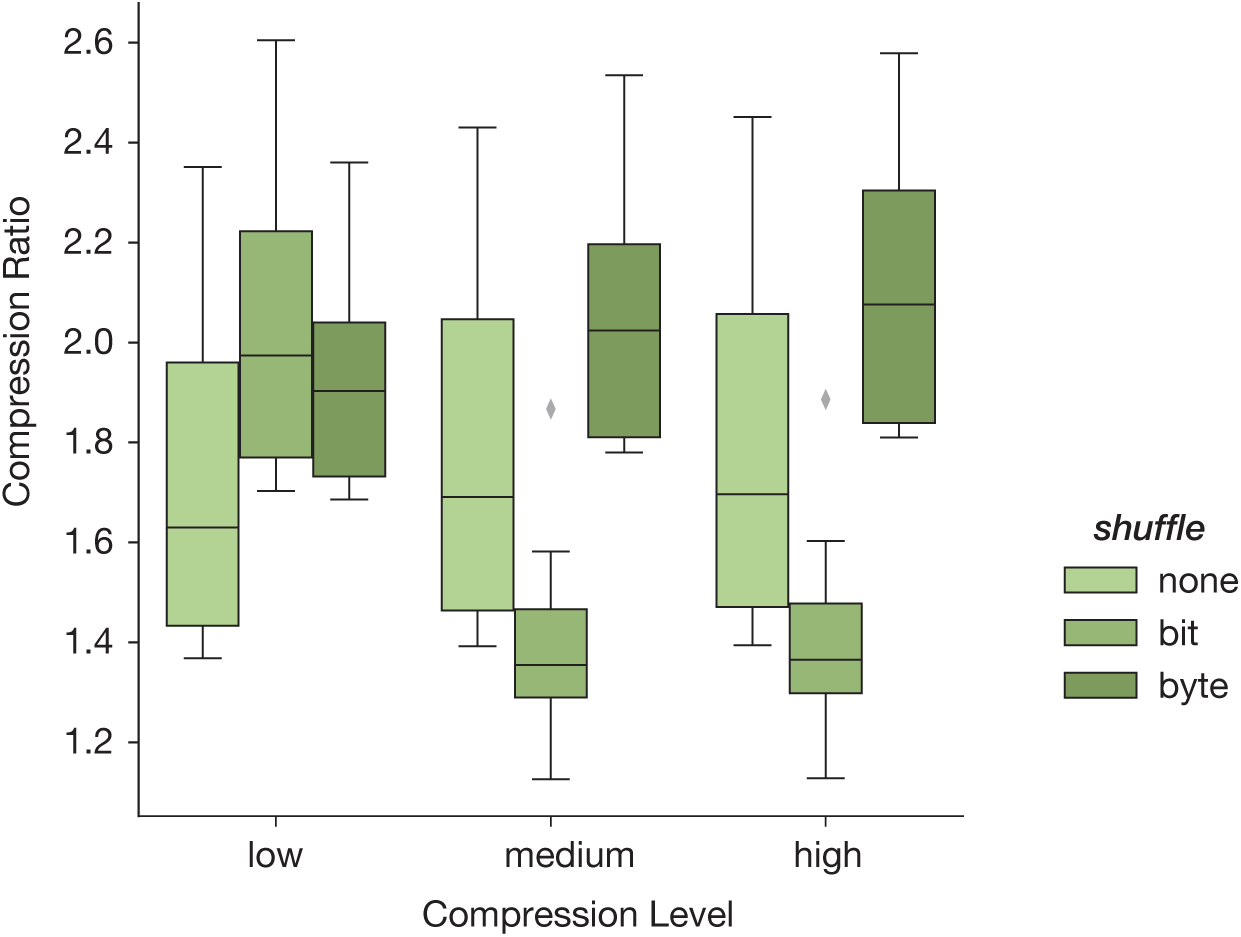
Combined effect of compression level and shuffling for blosc-zlib. Compression ratios for the `blosc-zlib` codec with increasing compression levels (*low* to *high*) for different shuffling options (*none*, *bit*, and *byte*). While compression ratio increases with increasing compression levels for *none* and *byte* shuffling options, the opposite behavior is observed for *bit* shuffling. *N* = 16 compression jobs for all distributions.

**Figure S2:**
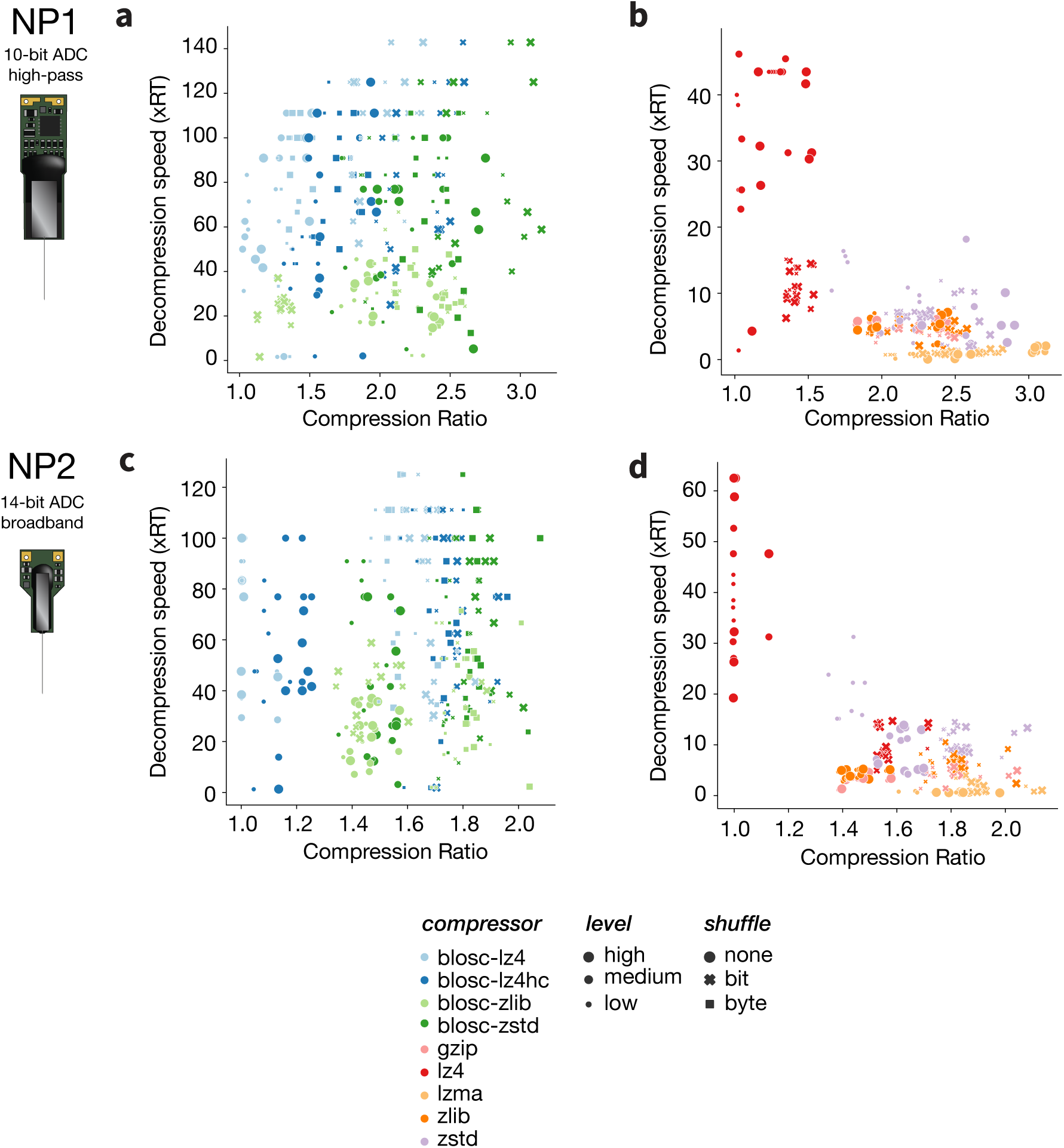
Compression ratio and decompression speed for all general-purpose compressors, shuffling options, and levels. Combined effect of compression level (marker size) and shuffling (marker shape) on compression ratio (*x*-axis) and decompression speed (*y*-axis) for *blosc* (**a** and **c**) and *numcodecs* (**b** and **d**) compressors. Top row is for NP1 (**a-b**), bottom row is for NP2 (**c-d**).

**Figure S3:**
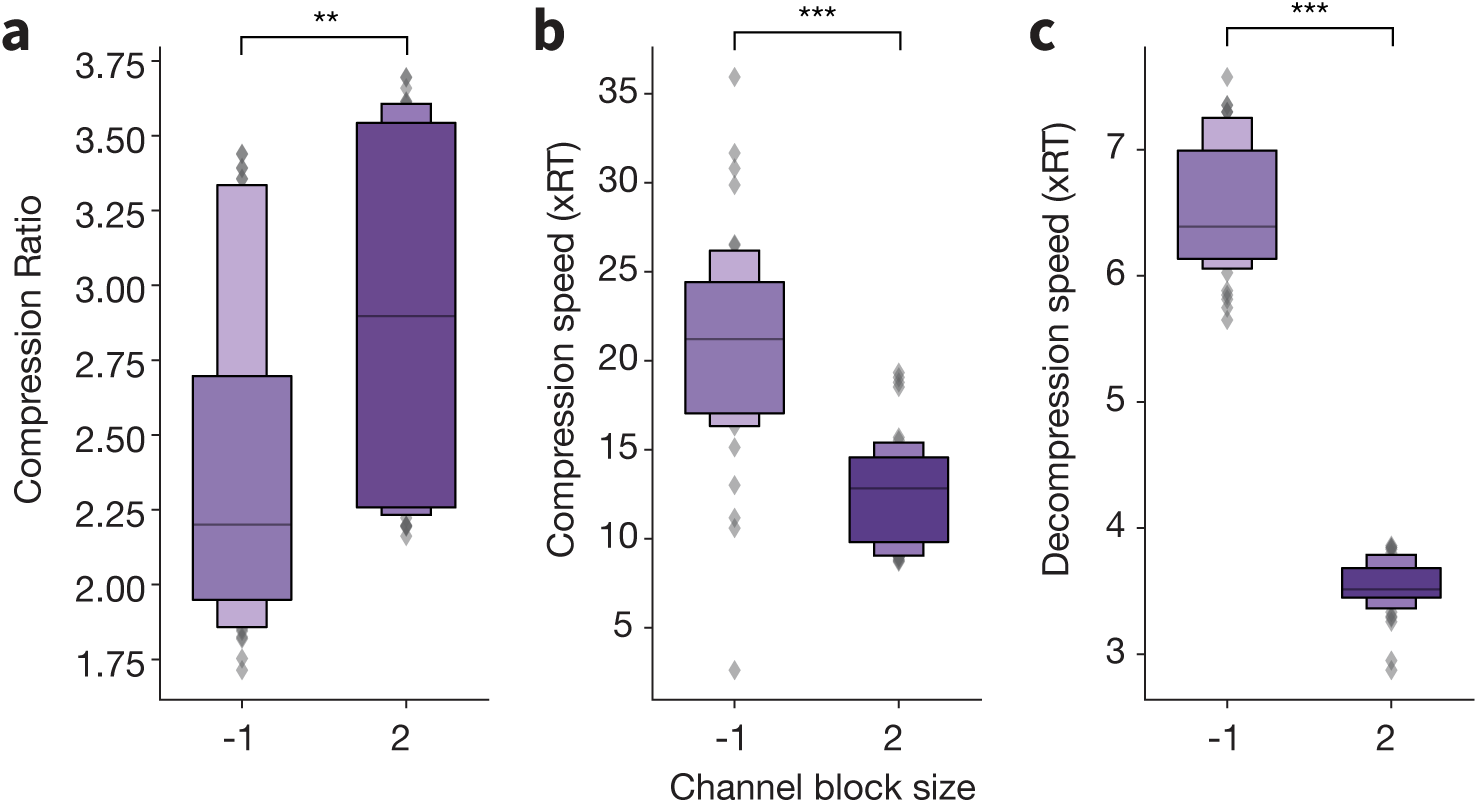
Effect of channel block size on FLAC compression. Compression ratio (**a**), compression speed (**b**), and decompression speed (**c**) for the `FLAC` codec with different channel block sizes. A channel block size of -1 corresponds to a flattening of the 2D input data before compression; a block size of 2 groups channels into *stereo* pairs. *N* = 48 compression jobs for all distributions.

**Figure S4:**
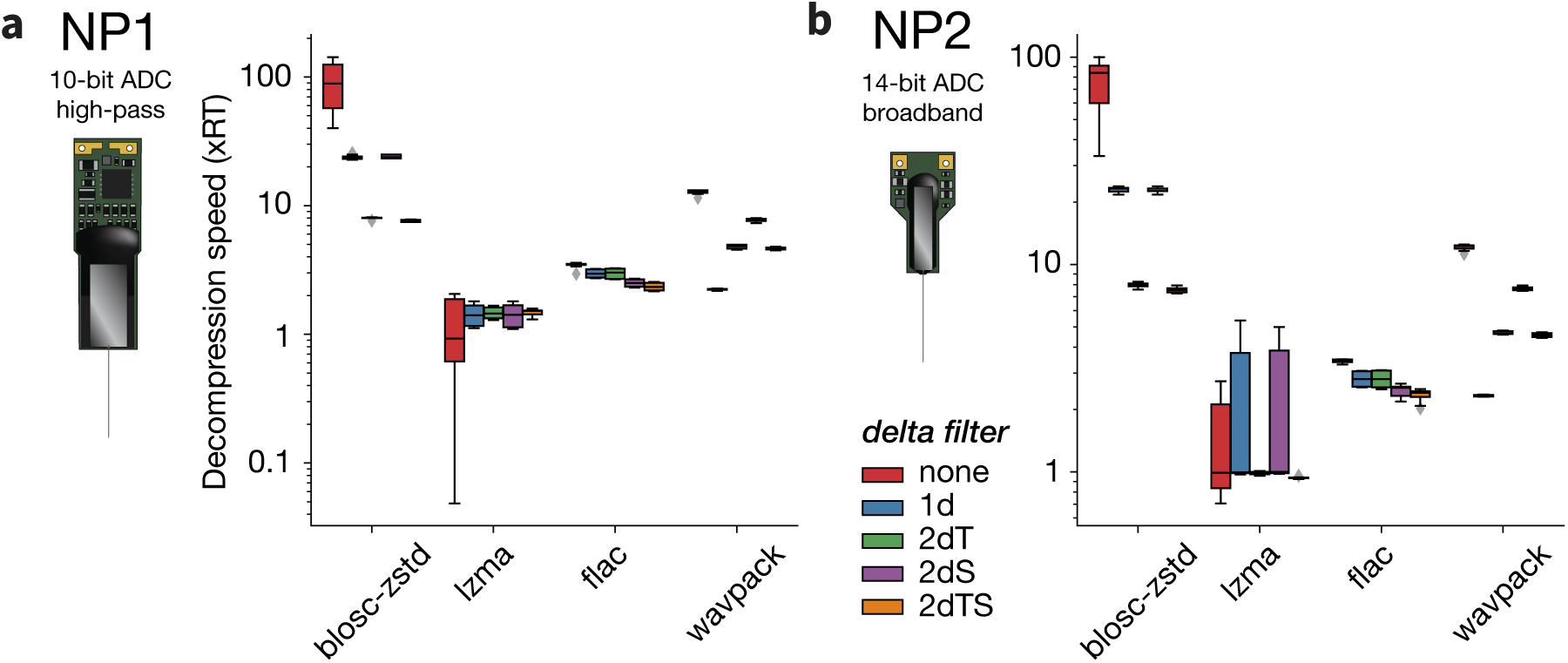
Effect of delta filters on decompression speed. Decompression speeds for `blosc-zstd`, `LZMA`, `FLAC`, and `WavPack` after applying different delta filters to NP1 (**a**) or NP2 (**b**) data. *N* = 8 compression jobs for all distributions.

**Figure S5:**
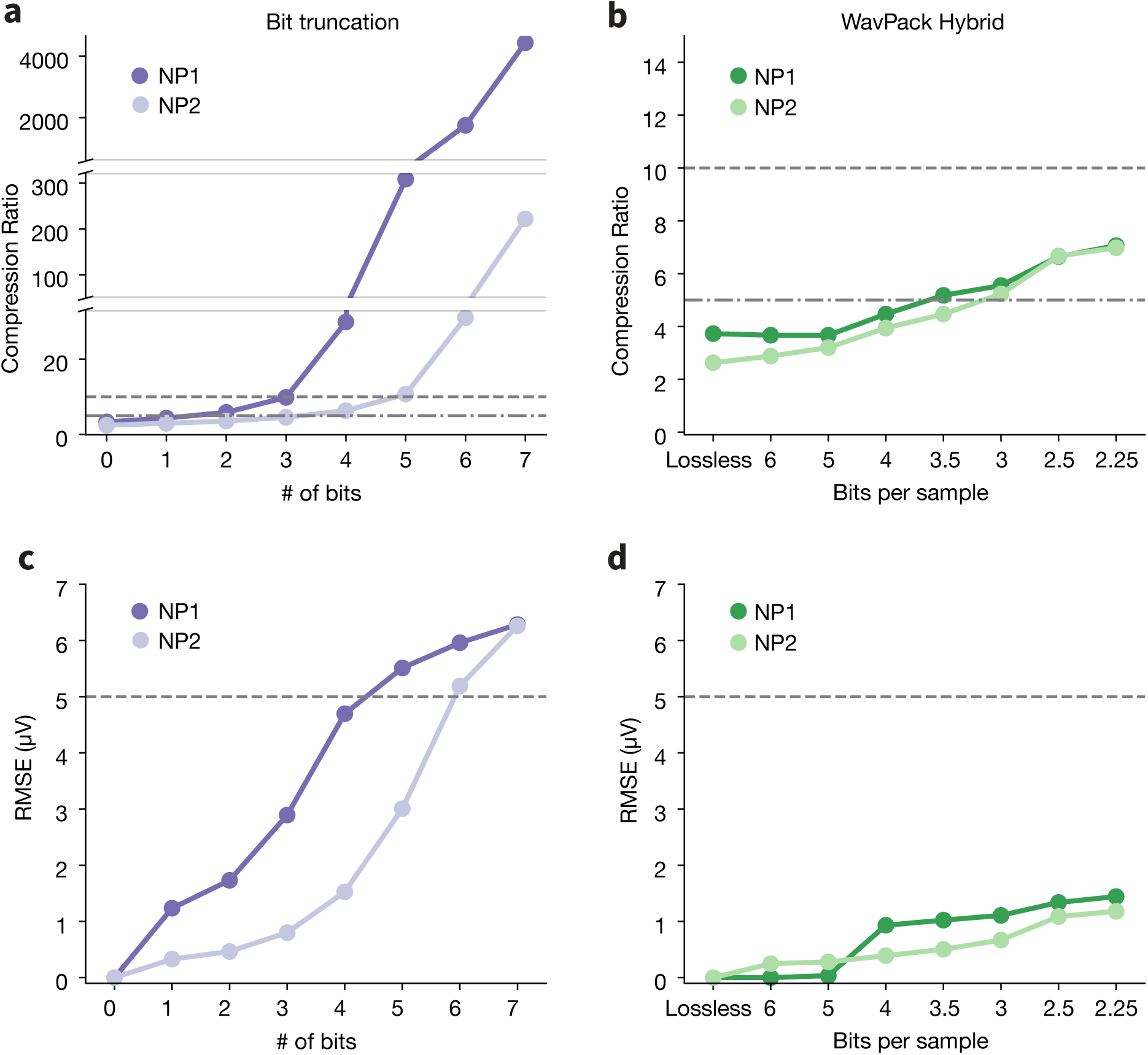
Impact of lossy compression on simulated data. Compression ratios (**a-b**) and root-mean-square error (RMSE) values (**c-d**) for bit truncation (left, purples) and `WavPack` Hybrid (right, greens). The *x*-axes represent the number of bits truncated (for bit truncation) or the target *bps* (for `WavPack` Hybrid). In all panels, lossiness increases from left to right. Horizontal dashed lines are added for reference and mark compression ratios of 5 and 10 or an RMSE of 5 µV.

**Figure S6:**
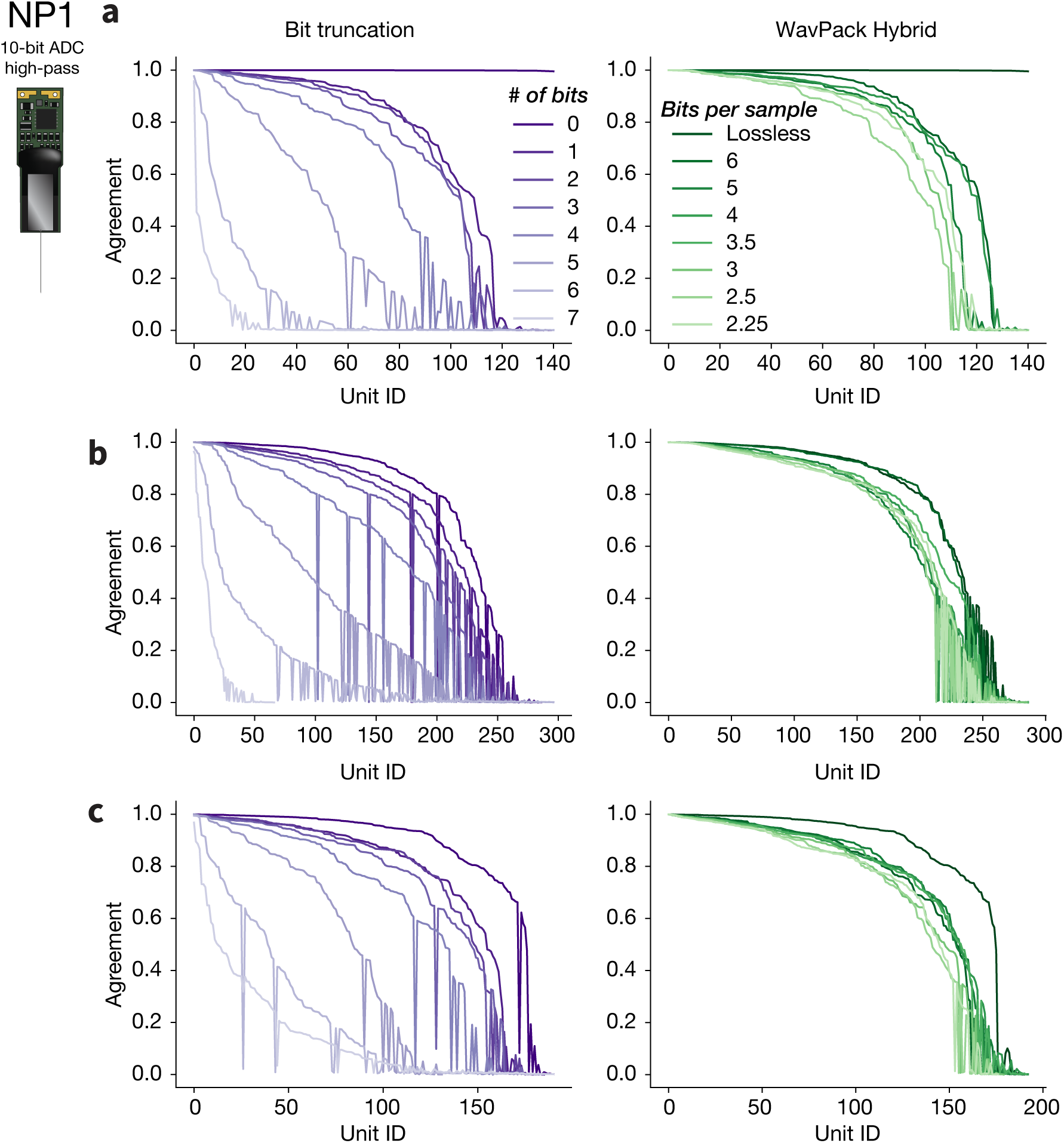
Comparing spike trains detected before and after lossy compression (NP1). Same as Figure 13 for the remaining NP1 sessions that were analyzed: **a**, 625749_2022-08-03_15-15-06_ProbeA. **b**, 634568_2022-08-05_15-59-46_ProbeA. **c**, SWC054_2020-10-05_probe00.

**Figure S7:**
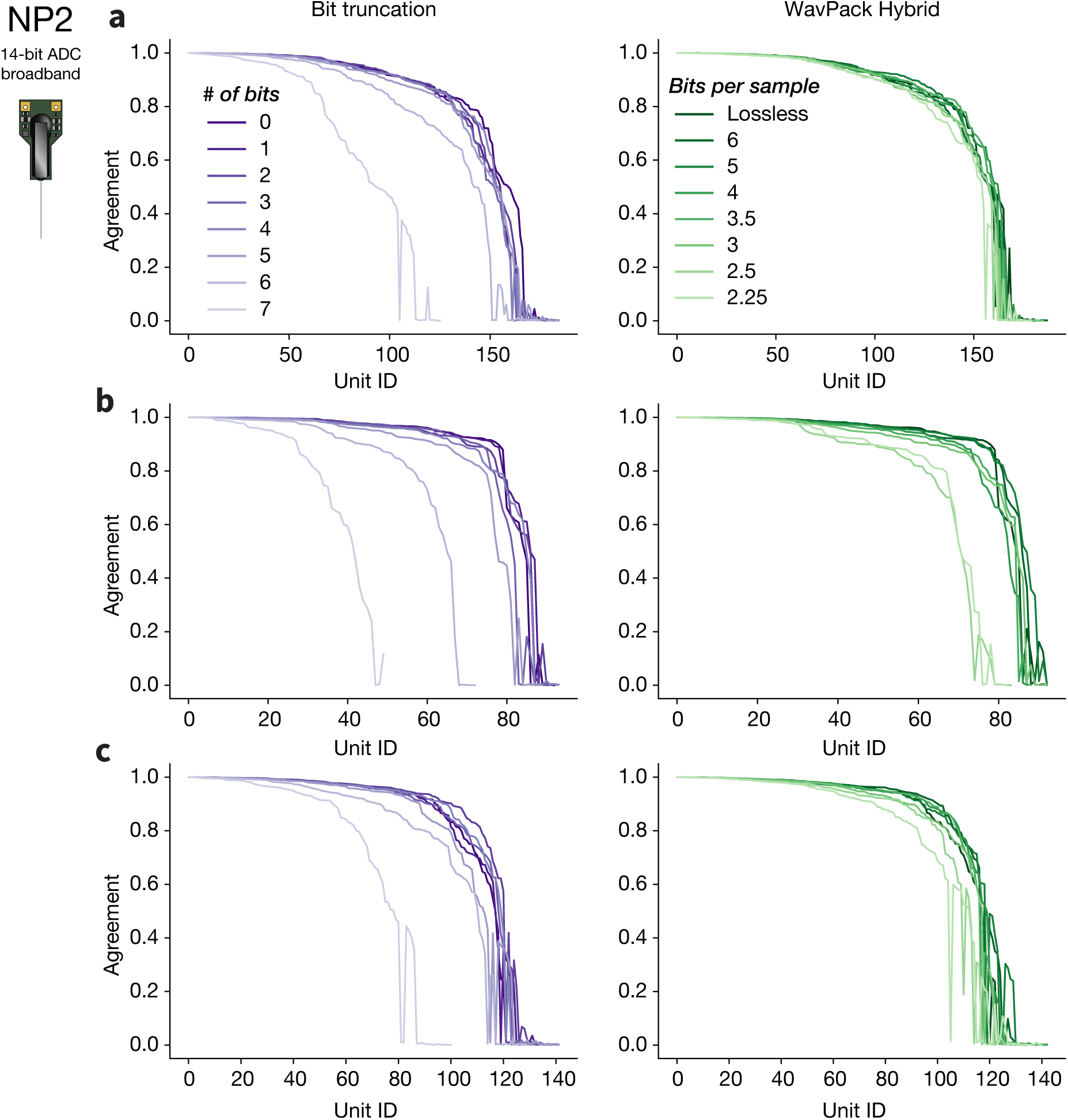
Comparing spike trains detected before and after lossy compression (NP2). Same as Figure 13 for the remaining NP2 sessions that were analyzed: **a**, 595262_2022-02-21_15-18-07_ProbeA. **b**, 602454_2022-03-22_16-30-03_ProbeB. **c**, 618384_2022-04-14_15-11-00_ProbeB.

**Figure S8:**
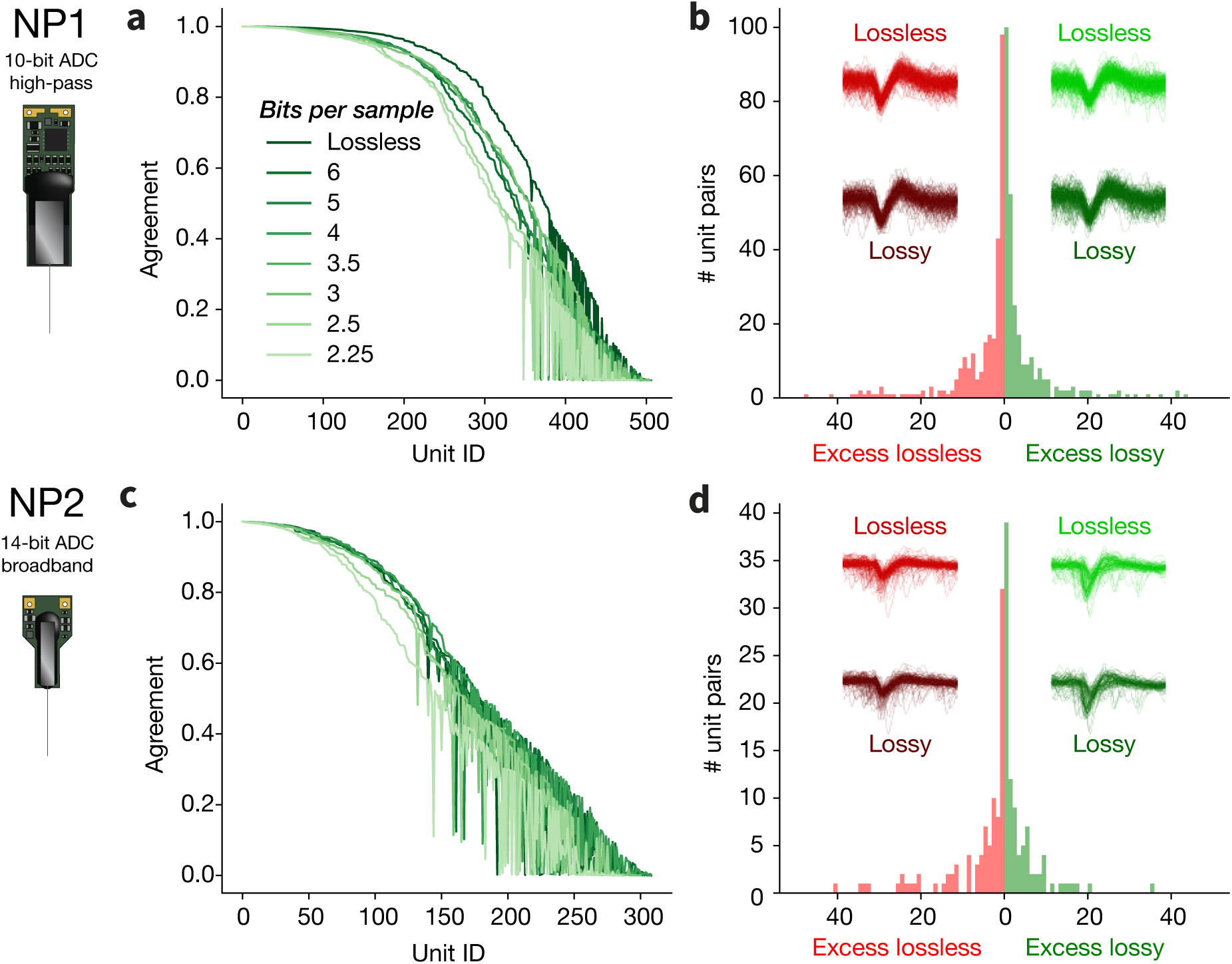
Extended comparison of lossless and lossy spike trains. Ordered agreement of all matching units between a lossless sorting output and seven levels of `WavPack` Hybrid compression, for NP1 (**a**) and NP2 (**c**) data (same datasets as Figure 13). For each matched unit with an agreement above 0.5, the number of excess spikes in the lossless and lossy (*bps* = 2.25) spike trains are shown (**b,d**). Excess spike waveforms for one example unit appear as insets: lighter waveforms are computed from the lossless recording, while darker ones are from the lossy recording.

## Footnotes

1 www.neuropixels.org

2 www.3brain.com/products/multiwell/hypercam-alpha

3 www.mxwbio.com/products/maxtwo-multiwell-microelectrode-array/

4 This calculation does not include the `LF` stream, which would result total rate of *∼*87 GB/hour.

5 as of March 2023, AWS: https://aws.amazon.com/s3/pricing/; GCP: https://cloud.google.com/storage/pricing

6 https://numcodecs.readthedocs.io/en/stable/

7 https://hdmf-zarr.readthedocs.io/en/latest/

8 https://github.com/int-brain-lab/mtscomp

9 https://medformat.org/

10 The final version of NP2 probes, which became publicly available in 2023, use 12-bit ADCs.

11 *Shuffling* refers to a re-organization of the samples before compression (at the byte or bit level) to make the data more compressible.

12 https://github.com/sonos/pyFLAC

13 Note that Neuropixels 2.0 are still at a prototype stage, and this value might differ from the final commercially distributed design.

14 A correction file can be generated to recover the original data, but we did not enable this option.

15 The *blosc* and `WavPack` codecs have inherently multi-threaded implementations, so some decompression steps did include parallelization.

16 **NP1**: 625749_2022-08-03_15-15-06_ProbeA, 634568_2022-08-05_15-59-46_ProbeA, CSHZAD026_2020-09-04_probe00, SWC054_2020-10-05_probe00; **NP2**: 595262_2022-02-21_15-18-07_ProbeA, 602454_2022-03-22_16-30-03_ProbeB, 612962_2022-04-13_19-18-04_ProbeB, 618384_2022-04-14_15-11-00_ProbeB – see Table 1

17 https://hdmf-zarr.readthedocs.io/en/latest/

18 https://portal.hdfgroup.org/display/support/Registered+Filter+Plugins

19 AWS on-demand pricing for an `m4` instance as of March 2023: https://aws.amazon.com/ec2/pricing/on-demand/

20 For simplicity, we assume that the decompression speeds can be achieved using a single CPU.

